# Method to determine whether sleep phenotypes are driven by endogenous circadian rhythms or environmental light by combining longitudinal data and personalised mathematical models

**DOI:** 10.1101/2023.06.14.544757

**Authors:** Anne C. Skeldon, Thalia Rodriguez Garcia, Sean F. Cleator, Ciro della Monica, Kiran K.G. Ravindran, Victoria L. Revell, Derk-Jan Dijk

## Abstract

Sleep timing varies between individuals and can be altered in mental and physical health conditions. Sleep and circadian sleep phenotypes, including circadian rhythm sleep-wake disorders, may be driven by endogenous physiological processes, exogeneous environmental light exposure along with social constraints and behavioural factors. Identifying the relative contributions of these driving factors to different phenotypes is essential for the design of personalised interventions.

The timing of the human sleep-wake cycle has been modelled as an interaction of a relaxation oscillator (the sleep homeostat), a stable limit cycle oscillator with a near 24-hour period (the circadian process), man-made light exposure and the natural light-dark cycle generated by the Earth’s rotation. However, these models have rarely been used to quantitatively describe sleep at the individual level. Here, we present a new Homeostatic-Circadian-Light model (HCL) which is simpler, more transparent and more computationally efficient than other available models and is designed to run using longitudinal sleep and light exposure data from wearable sensors. We carry out a systematic sensitivity analysis for all model parameters and discuss parameter identifiability.

We demonstrate that individual sleep phenotypes in each of 34 older participants (65-83y) can be described by feeding individual participant light exposure patterns into the model and fitting two parameters that capture individual average sleep duration and timing. The fitted parameters describe endogenous drivers of sleep phenotypes.

We then quantify exogenous drivers using a novel metric which encodes the circadian phase dependence of the response to light. Combining endogenous and exogeneous drivers better explains individual mean mid-sleep (adjusted R-squared 0.64) than either driver on its own (adjusted R-squared 0.08 and 0.17 respectively).

Critically, our model and analysis highlights that different people exhibiting the *same* sleep phenotype may have *different* driving factors and opens the door to personalised interventions to regularize sleep-wake timing that are readily implementable with current digital health technology.

**Author summary:** Disrupted sleep has long term health consequences and affects our day-to-day ability to function physically, mentally and emotionally. But what determines when and how long we sleep?

It is well-known that daily light exposure patterns determine the timing of the body clock. However, creating mathematical models that can take realistic light exposure patterns and predict plausible sleep timing has been challenging. Furthermore, nearly all previous studies have focused on developing models for average behaviour, yet sleep timing and duration are highly individual.

In this paper, we present a simple model that combines sleep regulatory and circadian processes. The model can take individual light exposure patterns and, by fitting physiologically plausible parameters, describe individual mean sleep timing and duration. We test our model on data collected from 34 older participants. Our modelling approach suggests that some of the participants slept late because of physiological factors, while for other individuals, late sleep was a consequence of their light environment.

This approach of combining a model with longitudinal data could be implemented in digital health technology such that your smart watch could tell you not only how you slept last night, but also how to change your light environment to sleep better tomorrow.

## Introduction

Sleep is a recurring physiological state in many species. In humans, it is a major determinant of quality of life, and disorders and disturbances of sleep are predictive of adverse health outcomes [1]. Sleep timing and duration change across the human life span but also show considerable individual differences in every age group. When individual differences in sleep timing are extreme, they are considered disorders; circadian rhythm sleep-wake disorders are prototypical examples [2]. Sleep disorders, including those associated with timing, are associated with psychiatric disorders and evidence indicates the relationship may be bidirectional [3, 4]. Interventions to ameliorate sleep timing are therefore desirable. Understanding, at an individual level, the endogenous versus exogeneous factors determining different sleep phenotypes [5] will facilitate the design of more effective, personalised treatments.

Sleep fulfils an essential need that accrues during wakefulness. Sleep homeostasis refers to the process that keeps track of this build up and dissipation of sleep need and this process is a determinant of sleep timing. Sleep timing in humans is also governed by a second physiological process, namely circadian rhythmicity. In the context of sleep regulation, circadian rhythmicity refers to the approximately 24-hour variation in sleep-wake propensity which persists even when sleep need and environmental factors such as light are kept constant. Circadian sleep propensity is high during the circadian night and low during the circadian day [6].

The alignment of circadian rhythms with respect to the 24-hour day is regulated by environmental light exposure patterns. The circadian pacemaker, located in the suprachiasmatic nuclei (SCN) in the brain, generates a near 24-hour rhythm with a near 24-hour period (24.15 h, standard deviation 0.2 h) [7]. This non-24-hour rhythm is synchronised to the 24-hour day by light input via a well-defined neural pathway connecting retinal photoreceptors to the SCN. The effects of light on the pacemaker depend on the phase of the circadian cycle at which light information reaches the SCN. Light in the biological morning speeds up the clock and light in the biological evening slows down the clock [8, 9]. Entrainment of our non-24-hour clocks to the 24-hour day occurs when there is an appropriate balance between ‘speeding up’ and ‘slowing down’.

Environmental light exposure patterns were once primarily driven by the natural light-dark cycle. However, in industrialised societies, most people spend approximately 90% of their time indoors [10] and, access to electric light means light is available at any time of day or night. Since indoor light is typically of lower intensity than outdoor light and we self-select when to turn lights on and off, the strength of the light-dark signal as a signal that can entrain the biological clock to the 24-hour day has been greatly degraded. The consequence of a less robust 24-hour light-dark signal is that for most people, i.e. those with an intrinsic period longer than 24-hours (77%), circadian rhythms and preferred sleep timing delay [11–14]. Social factors, such as work, school schedules and socialising, also influence sleep timing [15]. In sum, sleep homeostasis, circadian rhythmicity, the light-dark cycle, and social constraints interact to determine the timing of wake and sleep across the 24-hour day.

Understanding of the genetic molecular determinants and correlates of the sleep homeostat, circadian rhythmicity and the effects of light has grown rapidly in the past few decades (e.g. [16, 17]). At the same time laboratory studies have indicated that long and short sleepers differ with respect to the average level of homeostatic sleep pressure [18] whereas early and late sleepers differ in their intrinsic circadian period [19, 20]. However, at the individual level, understanding how these different regulatory mechanisms result in different sleep phenotypes in daily life is more limited. The difficulty of ascertaining causal factors driving sleep phenotypes is exacerbated by the fact that the same phenotype may have different underlying causes. For example, late sleep timing may be a consequence of too little bright light in the morning or too much bright light in the evening or a long intrinsic period (slow clock), or a combination of all of these. Parameters of the clock and sleep homeostasis can be quantified in laboratory protocols but these are labour intensive and are not scalable.

Mathematical models are one way to unravel the relative contributions of different factors. Several system-level mathematical models for the human sleep-wake cycle have been developed. Kronauer and colleagues modelled the sleep-wake cycle as an interaction between a strong circadian oscillator (driving core body temperature) and a weak sleep-wake oscillator. Entrainment to the 24-hour day occurred via forcing the strong oscillator by the effects of a so-called zeitgeber (‘time giver’) which entered the system via the weak oscillator [21]. However, the zeitgeber was not based on available data for the effects of light on the human circadian timing system. Daan, Beersma and Borbély modelled the interaction of the homeostatic and circadian process (hence the so-called ‘two-process’ model) but did not model the effects of light on the circadian process and hence could not simulate entrainment to the 24-hour day. Parameters of the homeostatic process of the two-process model were based on electroencephalography (EEG) slow wave activity (SWA, EEG power density in the 0.75-4.5 Hz range) and it was assumed that this variable played a key role in sleep timing. Parameters for the circadian process were motivated by average sleep duration following displacement of sleep with respect to the day by extending the wake period [22, 23].

The two-process model is a phenomenological model. Building on insights on the neuronal regulatory mechanisms and the conceptual flip-flop model [24], Phillips and Robinson developed a more physiologically-based model by considering mutual inhibitory sleep and wake promoting nuclei whose firing rates were modulated by circadian and homeostatic processes [25]. Later, they incorporated Kronauer’s model for the phase-shifting effects of light in phase response experiments [26, 27], by including a self-sustained oscillator and light input [28]. Previously, we have shown that an adapted version of the Phillips-Robinson model can accurately model sleep duration and timing across the lifespan [29] and have systematically investigated the impact of light and imposed social constraints (such as getting up for work) [14]. We have shown that combining light exposure data, collected longitudinally, with the Phillips-Robinson model can replicate average sleep timing and sleep duration within people living with schizophrenia and healthy unemployed individuals [30]. Using a similar model, Postnova and co-workers have modelled the effect of imposed wake schedules due to shiftworking [31] and in forced desynchrony experiments [32], on sleepiness, cognitive performance and ‘circadian dynamics’. Hong et al. have introduced the notion of ‘circadian sufficient sleep’ to quantify the degree of alignment of sleep and circadian rhythms [33]. In a further extension of the Phillips-Robinson model, Song et al. [34] predicted sleepiness based on individual sleep-wake history and proposed the duration and timing of the main sleep and a prophylatic nap to minimise sleepiness during shifts.

Concurrently neuronal models capturing wake, rapid eye movement (REM) sleep and non-REM sleep have been developed [35] which, consistent with observations, have modelled the differences between long and short sleepers by differences in mean levels of a sleep homeostat [36].

Neuronal models are specifically designed to incorporate more physiological insight into mathematical models of sleep-wake regulation than the phenomenological two-process model. However, consequently, they are more mathematically complex and have more parameters than are needed to explain currently available data on sleep timing and duration.

Here, we take a different approach and present a new, more minimalist, model that incorporates the effects of sleep homeostasis, circadian rhythmicity and light on sleep timing, which has a lower compute time and where the role of each parameter is more transparent. This new model has the intuitive appeal of the two-process model but also includes the effect of light on circadian timing. We undertake a systematic sensitivity analysis to find which parameters primarily regulate sleep timing and duration and discuss which parameters are identifiable from light and sleep timing data. In older participants (N=34, age 65-83 y), we demonstrate that we can extract individual parameters describing endogenous physiological processes by combining the model with data from wearable light sensors and fitting for individual average sleep duration and timing. Associations between model parameters and multiple sleep measures, including standard biomarkers of sleep homeostasis collected in the laboratory, are considered. The exogenous driver of light is quantified using both standard metrics of light exposure and a novel light metric that quantifies both the intensity and time-of-day dependent effects of light on the biological clock. Finally, using the extracted parameters and the novel light metric, we propose a method to identify the endogenous physiological versus the exogenous environmental factors underlying different phenotypes and discuss the implications for the design of light interventions delivered by digital health technology applications.

## Methods

### Ethics statement

The study received a favourable opinion from the University of Surrey Ethics Committee (Approval number: UEC 2019 065 FHMS) and was conducted in accordance with the Declaration of Helsinki. Written informed consent was obtained from participants before any procedures were performed.

### A new mathematical model for realistic sleep timing

Our Homeostatic-Circadian-Light (HCL) model is motivated by elements of the original two-process model for sleep-wake regulation [22, 23], the neuronal model of Phillips-Robinson model [25, 28] and a model of the interaction of light with the human biological clock [27]. The model is illustrated in Fig 1 and described in detail below.

**Fig 1.**
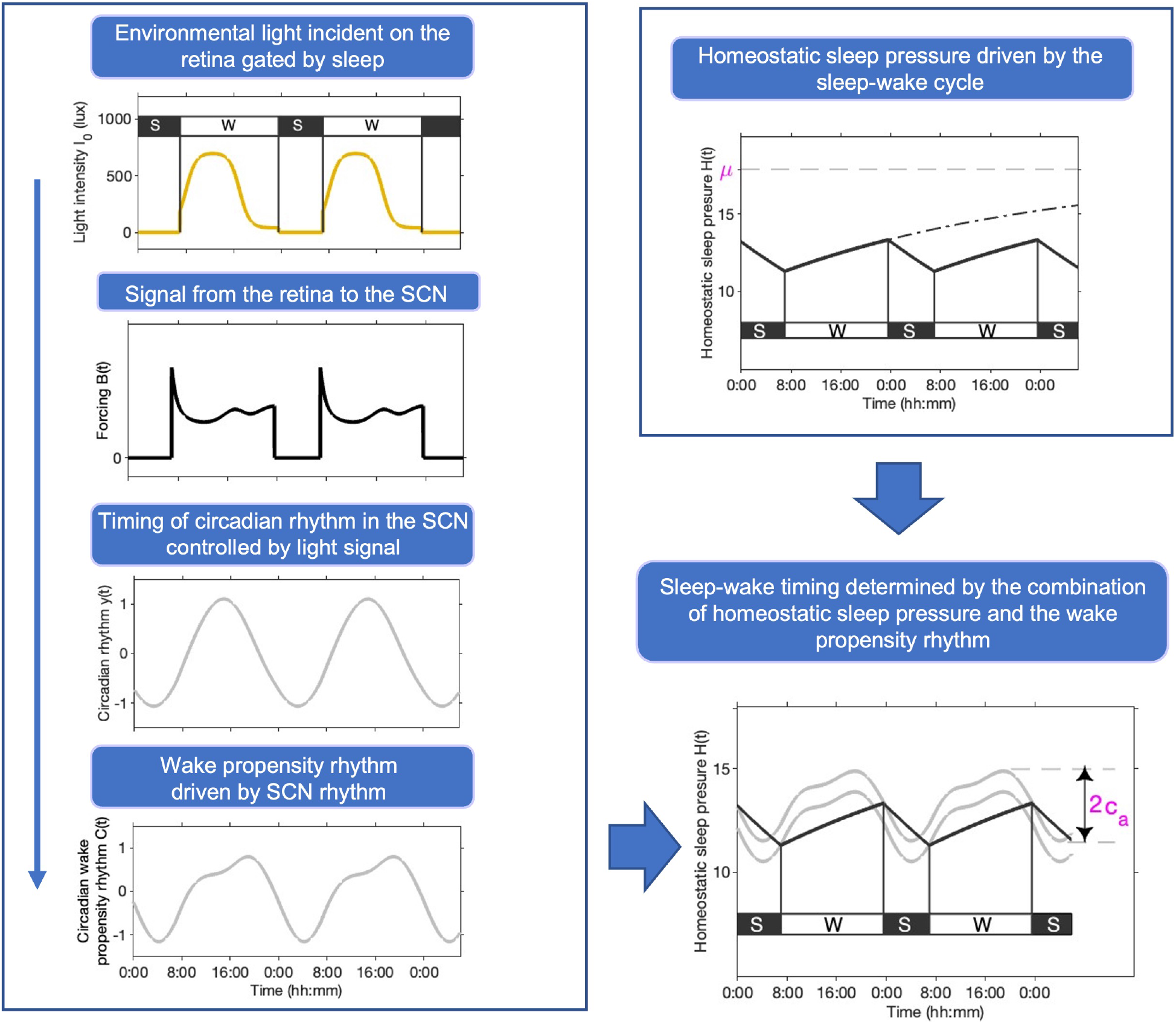
Homeostat-Circadian-Light (HCL) model. Patterns of available environmental light gated by the sleep-wake cycle (since we turn off the lights and close our eyes when we go to sleep), result in a rhythmic signal that passes from the retina to the suprachiasmatic nuclei (SCN). Provided the signal is of sufficient ‘strength’, it controls the timing of rhythms in the SCN, which in turn drives the 24-hour rhythm in the circadian drive for wakefulness and sleep. Homeostatic sleep pressure *H* increases during wake and decreases during sleep. If kept awake, homeostatic sleep pressure asymptotes to an asymptote *μ*. The larger the value of *μ*, the faster the rise rate of homeostatic sleep pressure during wake. Switching between wake (W) and sleep (S) states occurs at thresholds modulated by the circadian rhythm of wake-sleep propensity [37] which has peak-to-peak amplitude 2*c*_*a*_.

#### Sleep homeostasis

As in the two-process model, it is assumed that there is a sleep pressure signal *H*(*t*) which increases during wake and decreases exponentially during sleep. Specifically,

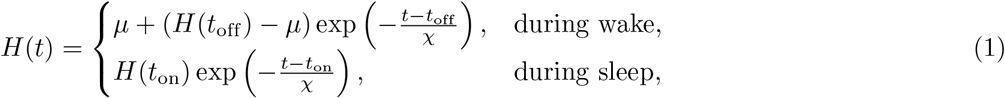

where *t* is time, *t*_on_ and *t*_off_ are the times of sleep onset and offset respectively, *χ* describes the rate of decay of sleep pressure during sleep whereas the rise rate during wake is governed by both *μ* and *χ*. The parameter *μ* is the ‘upper asymptote’ for the homeostatic sleep pressure signal since during wake *H*(*t*) → *μ* as *t* → ∞. In other words, if kept awake for a long time, homeostatic sleep pressure approaches an upper limit that cannot be surpassed, see Fig 1. Similarly, homeostatic sleep pressure approaches a lower asymptote of zero during sleep.

In differential equation form,

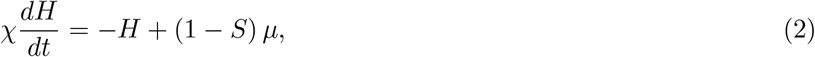

where *S* takes the value 1 during sleep and 0 during wake.

#### Circadian rhythmicity

As in the two-process model, we assume that spontaneous switching from wake to sleep and from sleep to wake occurs at thresholds which are periodically modulated to mimic the effect of circadian rhythmicity. Hence, switching from wake to sleep occurs at *H*(*t*) = *H*^+^(*t*) where

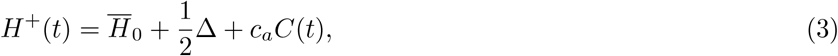

and switching from sleep to wake occurs at *H*(*t*) = *H*^−^(*t*) where

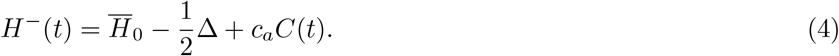

The constant 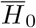 is the mean level of upper and lower thresholds and Δ is the separation between the thresholds.

The wave form of the time dependent circadian modulation *C*(*t*) represents the circadian rhythm of wake propensity. Evidence for a strong circadian drive for ‘wakefulness’ has been observed both for sleep measures [37] and for cognitive measures [38]. For the functional form used here we have fitted to experimental data from forced desynchrony experiments [37]. The functional form is discussed in greater detail in the subsection ‘Light coupling to circadian rhythmicity’ and in Section B in S1 Appendix. Paradoxically, but consistent with data and the understanding that sleep and circadian processes act in opposition, the circadian wake propensity signal increases during the ‘day’, reaching a maximum shortly before bedtime and a minimum in the second half of the night

Equations (2)-(4) may be considered as a lower dimensional representation of a neuronal model in which switching between wake and sleep occurs on a fast timescale. The neuronal interpretation of 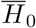 is that it represents the drive to be awake, for example from orexin [24], whereas Δ represents the strength of the mutual inhibition between sleep and wake promoting neurons. Further discussion of the link between 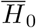 and Δ and parameters in neuronal models such as [28] is given in Section A in S1 Appendix.

#### Light coupling to circadian rhythmicity

The function *C*(*t*) describes the circadian wake propensity rhythms and has a peak-to-peak amplitude of approximately two. Here, the circadian rhythm *C*(*t*) is constructed from a model of the effect of light on circadian phase and data describing the circadian drive for wakefulness [37]. Specifically, following Kronauer [26], the circadian timing system is modelled as the interaction of Process L (L for light), and Process P (P for pacemaker) [26]. In Process L, light is considered to activate photoreceptors in the eye and result in a driving ‘force’, 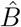, on the circadian pacemaker,

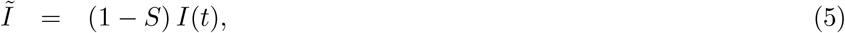

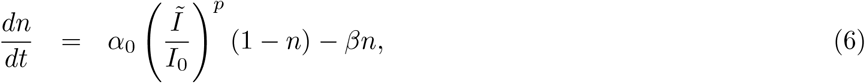

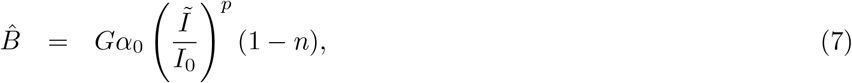

where *I*(*t*) is the available light, 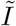 is the light that reaches the retina and *n* is the fraction of activated photoreceptors. The factor (1 − *S*) in equation (5) models the fact that eyes are closed during sleep, so that sleep has a gating effect on light exposure. Equation (6) states that when light reaches the retina, the rate of saturation of photoreceptors is given by 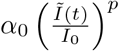 and the rate of decay of activated receptors is given by *β*. The values of *α*_0_, *I*_0_, *p* and *β* have been determined from experimental data collected during phase response experiments [26, 27]. Of particular note is that the response to light of intensity 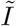 is ‘compressed’ since it appears in the form 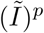 and *p <*1.

The pacemaker P is modelled as a van der Pol oscillator, here, as in [27], given by,

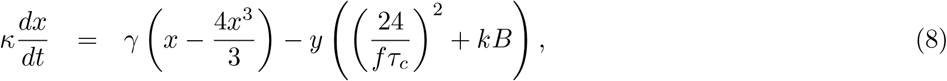

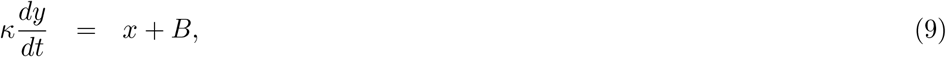

where 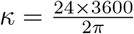 is a time scaling factor and

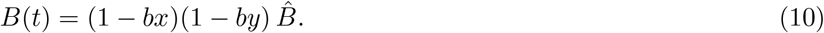

A typical form for *B*(*t*) for one specific light profile is shown in Fig 1 and shows that it incorporates both so-called ‘phasic’ and ‘tonic’ elements [39]. Phasic effects induce abrupt changes in the position of the oscillator, as occurs as a result of the spike in *B*(*t*) at the transition from dark to light. Tonic effects change the speed of the clock through modulating the period of the pacemaker and occur via the fact that *B*(*t*) settles to an approximately constant value in constant light.

In equations (8), (9), *y*(*t*) represents the circadian pacemaker activity, and *x*(*t*) is an auxiliary variable. The physiologically motivated parameters result in the van der Pol oscillator operating in its weakly nonlinear regime where *x*(*t*) and *y*(*t*) oscillate in a close to sinusoidal manner, see Fig 1. In the absence of light, these oscillations have a natural period, the ‘intrinsic circadian period’, of *τ*_*c*_ hours. The parameter *γ* is the stiffness of the oscillator which describes how easily the system is perturbed from its natural oscillatory state and the speed of recovery following perturbation. Consistent with observations that to induce large changes in phase in humans repeated bouts of high intensity light are required [40], the value of *γ* is taken to be relatively large. This means that the periodic solution (limit cycle) is strongly attracting,

The effect of light on the van der Pol oscillator comes from the light drive, 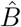 modulated by a sensitivity function (1 − *bx*)(1 − *by*) that describes how the strength of the drive varies with the position of the oscillator.

In the HCL model, the function *C*(*t*) is taken to be

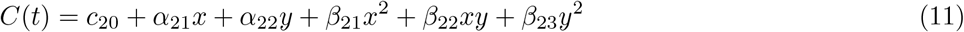

where the coefficients *c*_20_, *α*_*ij*_ and *β*_*ij*_ were found by fitting to the circadian drive for wakefulness reported in [37], see Fig 1. Further details of the fitting process are given in Section B in S1 Appendix, along with the values of the coefficients.

Hence the complete HCL model consists of four differential equations: one for the sleep homeostat, (2); two for the van der Pol oscillator, (8), (9), and one for the photoreceptors, (6). In addition, there are the two switching conditions: switching from wake to sleep occurs when *H*(*t*) = *H*^+^(*t*) and switching from sleep to wake when *H*(*t*) = *H*^−^(*t*), where *H*^*±*^(*t*) are given by equations (3), (4) and (11). Finally, the light exposure pattern *I*(*t*) is needed as an input, see equation (5). In this paper, *I*(*t*) will come from light intensity data measured in individuals. For scenario testing and intervention design, synthetic profiles are also used.

#### Parameter values in the HCL model

The HCL model contains a total of 14 parameters, five directly linked to the sleep-wake regulation aspects of the model 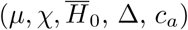 and nine related to the interaction of light with the pacemaker (*fτ*_*c*_, *G, p, k, B, γ, α*_0_, *β, I*_0_).

Default parameter values for *χ*, 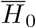 and Δ are motivated by considering the HCL model as a slow manifold reduction of the neuronal model in [14]. Hence the gap between the thresholds Δ is much smaller, and typical values for the circadian amplitude *c*_*a*_ are much larger, than those used in the original two-process model. Furthermore, consistent with neuronal models, we take the homeostatic time constant *χ* to be 45 hours whereas the two-process model has different time constants for the rate of rise of sleep homeostasis during wake and rate of fall of sleep homeostasis during sleep. These time constants have been measured from slow wave activity in sleep electroencephalography (EEG) studies to be approximately 18 hours and four hours respectively [23]. The HCL time constants appear very different, but since the actual rise / decay rates of the homeostat depend not only on the homeostatic time constants but also on the values of the upper (and lower) asymptotes it is not easy to make a direct comparison. Further comparison between the HCL model parameter regime and that of the original two-process model is given in Section C and Fig B in S1 Appendix. In S1 Appendix Section C we also demonstrate that the ratio of the average rate of rise of sleep homeostasis during wake compared with the average fall during sleep is given by the ratio of sleep to wake duration). Thus for an average sleeper who is asleep for 8 hours and awake for 16 hours, the ratio is one half.

For the purposes of this paper, default values for *μ* and *c*_*a*_ were chosen such that, with the same default light profile used in [14] and shown gated by sleep in Fig 1, mid-sleep time was 03:16 hh:mm. This matches mid-sleep time data reported in [41, 42] for people aged approximately 65 y. Default values for the ten light / circadian parameters were fixed to the standard parameters used in [14, 28, 30].

All parameters, their meanings and their default values are listed in Table 1.

**Table 1.** Default parameter values for the HCL model and their meanings.

| Sleep / wake regulation parameters |  |  |
| --- | --- | --- |
| Parameter | Value | Meaning |
| $\mu$ | 17.87 | Upper asymptote giving the upper bound for homeostatic sleep pressure and controlling the rate of rise of homeostatic sleep pressure during wake. |
| $\chi$ | $45 \times 60 \times 60$ s | Time constant for rise and decay of homeostatic sleep pressure. |
| $\bar{H}_0$ | 13 | Mean level of the wake propensity rhythm. |
| $\Delta$ | 1 | Separation between the thresholds. |
| $c_a$ | 1.72 | Circadian amplitude. |
| Light / circadian parameters |  |  |
| Parameter | Value | Meaning |
| $\tau_c$ | 24.2 h | Circadian period. Natural period of the van der Pol oscillator. |
| $f$ | 0.99669 | Correction factor so that in the absence of light the van der Pol oscillator gives oscillations with a period of $\tau_c$. |
| $G$ | 19.9 | Gain factor determining the magnitude of the effect of light on the driving force $\hat{B}$ on the circadian pacemaker. |
| $p$ | 0.6 | Light sensitivity. Determines the steepness of the dose response curve to light. |
| $k$ | 0.55 | Determines the relative effect of light on the oscillator variables $x$ and $y$. It changes the shape of the velocity response curve. |
| $b$ | 0.4 | Sensitivity modulation factor that determines the degree to which the strength of the drive on pacemaker P depends on the position of the pacemaker. Shifts the time of maximum sensitivity to light. |
| $\gamma$ | 0.23 | Stiffness of the van der Pol oscillator determining how quickly oscillations return to the limit cycle when perturbed. |
| $\alpha_0$ | $0.16/60$ s<sup>-1</sup> | Factor determining the magnitude of the effect of light on the fraction of activated photoreceptors and on the drive. |
| $\beta$ | $0.013/60$ s<sup>-1</sup> | Decay rate of fraction of activated photo receptors. |
| $I_0$ | 9500 lux | Scaling factor for light. |

#### Differential equation solver

Simulations of the HCL model were carried out in MATLAB [43] using the stiff solver ode15s with relative and absolute tolerances set to 10^−4^. The event solver was used to locate switching points from wake to sleep and from sleep to wake.

### Data collected in everyday life

The primary purpose of developing the HCL mathematical model was to construct a simpler, more transparent model that was computationally less complex than existing models and could be used to fit longitudinal data for deeper understanding of causal factors driving sleep phenotypes in close to real time.

In order to test our approach, we used light, actigraphy and sleep diary data collected for 7-14 days at home from 35 participants (age (mean ± SD) 70.9 ± 5 years, range 65–83 years, 14 women). Immediately after the 7-14 days at home, participants undertook an overnight polysomnography (PSG) recording at the Surrey Sleep Research Centre, Guildford, UK.

To be deemed eligible for the study participants had to be between the age of 65 and 85 years, be self-declared mentally stable and physically healthy but could have stable controlled medical conditions including Type 2 diabetes, hypertension, arthritis. They needed to be able to perform daily living activities independently, drink less than 29 units of alcohol per week and be non-smokers. Eligibility for the study was assessed via a telephone interview and an in-person screening visit which included provision of demographic information, measurement of height, weight and vital signs, self-reported medical history, symptom directed physical examination and completion of a series questionnaires including the Epworth Sleepiness Scale [44] and the Pittsburgh Sleep Quality Index [45]. Data were collected from the first 18 participants from January to March 2020. Following a temporary halt in data collection due to COVID, data from the remaining 17 participants were collected from July to November 2021.

Subjective sleep-wake patterns were assessed via daily completion of the Consensus Sleep Diary [46]. Parameters collected through the sleep diary included the time the participant went to bed, when they tried to go to sleep, how long it took them to fall asleep (sleep latency), number of awakenings, and time of final awakening. For the purposes of this paper, sleep onset times for each participant were calculated as the sleep diary time when participants started to try to go to sleep plus the sleep latency (i.e. answers to the questions: What time did you try to go to sleep? How long did it take you to fall asleep?). Sleep offset times were considered to be the sleep diary final awakening time (i.e. answer to the question: What time was your final awakening?).

Light exposure patterns were assessed by continuous wearing of a wrist worn actiwatch (Actiwatch Spectrum, Philips Respironics, Murrysville, PA, USA). Environmental white light levels, recorded in lux at one minute intervals, were extracted from the Actiwatch datasets. One participant was excluded from the fitting because the Actiwatch stopped recording after two days. The second group of participants in addition wore a second light measuring device (HOBO Pendant MX2202 Onset Data Logger, Bourne, MA, USA) that was worn as a pendant and clipped to clothing near the shoulder. Results were calculated both using data recorded at the wrist and data recorded at the shoulder to test the sensitivity of results to the position of the sensor.

Since participants wore an Actiwatch spectrum an alternative to using the participant self-reported sleep diaries would have been to use outputs from the Actiwatch algorithms for automated analysis of rest-activity records. However, for the participants reported here, we have shown that sleep diaries more accurately assess nocturnal sleep-timing parameters than Actiwatch outputs that are unassisted by sleep diaries to set the analysis period [47].

#### Light imputation

Participants were encouraged to keep all light monitors uncovered. Nevertheless, there were some periods in which the participants reported being awake but the recorded lux values were zero. For values that were zero and occurred during wake, imputation was carried out and zero values were replaced by the median value for the half an hour before and half an hour after, calculated across all available days of data for the participant concerned. Fittings were carried out with both raw and imputed light data to test the sensitivity of the method to imputation.

Light was recorded at one minute intervals. When the ordinary differential equation solver used to solve the HCL equations required values of light at intermediate intervals, light data were linearly interpolated.

#### Slow wave activity

Full PSG data were collected using the SomnoHD system (SOMNOmedics GmbHTM, Germany) according to the American Academy of Sleep Medicine (AASM) guidelines during the overnight in-lab recording which consisted of an in-bed period of ten hours. The Domino software associated with the SomnoHD system was used to score sleep at 30 s intervals [48]. A consensus hypnogram was generated using the manual sleep scoring performed by a registered polysomnographic technologist and an experienced scorer. The six channels of electroencephalography (EEG) data collected in the PSG included frontal (F3-M2; F3-M1), central (C3-M2; C4-M1) and occipital (O1-M2; O2-M1) derivations. The artefacts in the EEG were manually scored by an expert and the labels were verified by a second expert to ensure accurate removal of the artefacts.

Slow wave activity (SWA) was calculated according to the method outlined in [49]. Specifically, each 30 s epoch that was classified as sleep was further divided into 4 s sub-epochs with an over-lap of 1s (10 sub-epochs per 30 s epoch). The artefacts in each of the 4 s sub-epochs were manually scored by an expert and the labels were verified by a second expert to ensure accurate removal of the artefacts. Power spectral density estimates were created via Fast Fourier Transform (FFT) for each of the artefact free 4 s sub-epochs and a measure of SWA activity for each 30 s epoch was calculated by averaging power in the 0.75-4.5 Hz range [50]. The timeseries of SWA for the first 3 hours of the recording from the two frontal and two central derivations were averaged to produce a single measure of SWA for each participant.

#### Parameter fitting

For each participant, the light data were fed into the HCL model and then parameters were found that minimized the difference between the average diary sleep duration and the average modelled predicted sleep duration and the average diary mid-sleep timing and the average model predicted mid-sleep timing. Care was taken to remove transients in the calculation of sleep duration and mid-sleep time by starting from default settings for the initial conditions for *H, x, y* and *n* and then integrating repeatedly for the duration of the light data (truncated to an integer number of days) until the model predicted average sleep duration and mid-sleep time had converged.

In order to test the robustness of our modelling approach, sleep data from each participant were fitted four times for the first cohort in which light data were collected at the wrist, and eight times for the second cohort in which light data were collected from both the wrist and shoulder. First, the impact of light imputation was tested by fitting using both raw and imputed light data. Next, the impact of the position of the sensor was tested by fitting using both light data collected from the shoulder and from the wrist. In all cases, fitting was carried out for the pair of parameters (*μ, τ*_*c*_) and separately for the pair of parameters (*μ, c*_*a*_) to investigate different possible physiological interpretations. The motivations for choosing these two pairs of parameters are discussed in the results.

The fittings were implemented in MATLAB. For reasons of computational speed, different minimization methods were chosen depending on the fitting parameters. Specifically, when fitting for the pair of parameters (*μ, c*_*a*_) the sum of squared residuals between the observed and model predicted averages of sleep duration and mid-sleep time was minimized using a Nelder-Mead search algorithm (fminsearch). The optimization routine was considered to have successfully converged to a solution if the residuals were less than 0.03 and the parameters were physiologically realistic. In particular, we required all parameters to be positive and 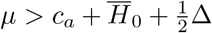. The reason for imposing the latter constraint is that when 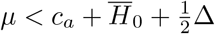, there are some circadian phases close to the maximum wake propensity at which it is impossible for homeostatic sleep pressure to reach the upper threshold and therefore for the model to spontaneously fall asleep. At first sight, this seems in line with the literature where the region close to the maximum wake propensity is known as the wake maintenance zone or forbidden zone for sleep and is a region where sleep is less likely to be initiated [51]. In fact, the presence of the wake maintenance zone in the model follows from the oscillatory shape of the threshold. The reason for judging values of *μ* that violate the imposed constraint as ‘non-physiological’ is because they lead to the model predicting that, even if someone had been kept awake for days, spontaneous sleep could not be initiated at these circadian phases.

Fitting for the pair of parameters (*μ, τ*_*c*_) was carried out iteratively using knowledge of model behaviour, as reported in the results. Specifically, sleep duration is almost independent of *τ*_*c*_ and depends monotonically on *μ*, whereas sleep timing depends monotonically on *τ*_*c*_, see Section. Hence, first the value of *μ* that gave a model predicted average sleep duration that matched the average diary sleep duration was found using a bisection-based method (fzero). Second, the value of *τ*_*c*_ that matched the average diary mid-sleep time was found using a bisection-based method (fzero). The process was repeated if the sum of the squared residuals between the observed and model predicted averages of sleep duration and mid-sleep time did not meet the tolerances specified. In both cases, an interval containing the solution was first found based on the known monotonic dependence of the outcome measure on the parameters. Fitting iteratively was faster and more reliable than using fminsearch.

### Metrics for light exposure patterns

Light exposure patterns can be quantified in different ways, and it remains unclear which metric is most relevant for light’s effect on the circadian system. Here, five standard metrics were calculated for each participant for each complete day of data: the number of hours of bright light exposure (light *>* 500 lux); the mean lux value; the geometric mean light exposure calculated by first taking the mean of the log (lux+1) and then raising the resulting value to the power of 10; the mean time of day at which half the daily light exposure occurred by considering the time at which half the cumulative daily lux occurred; the mean time of day at which half the daily light exposure occurred by considering the time at which half the cumulative daily log(lux+1) occurred. These metrics are illustrated in Fig C in S1 Appendix.

#### Novel light metric quantifying the biological effect of light

In addition, a novel measure of the biological effect of light was constructed by feeding each day of light data into the light-circadian model of Forger et al [27] (essentially equations (5 - 10) for default parameters and *S* = 0) and integrating for one day. Initial conditions were set to be those for the state of the light-clock system (given by *x, y* and *n*) at midnight for someone with a mid-sleep of 04:30, a sleep duration of 8.22 h and the synthetic light profile used in [14]. These values were chosen to be consistent with the ‘average’ person considered in [14]. The light-circadian model is designed to capture the interaction of light on the human velocity response curve, including the fact that at some times of day light speeds up the clock and at others it slows down the clock. Integrating for one day and considering whether the clock has covered more than or less than one cycle (here expressed in minutes) quantifies the extent to which that day of light data has a net effect of speeding up or slowing down the clock. Positive values mean that the clock has undergone more than one cycle in 24 hours, negative numbers mean that the clock has undergone less than one cycle in 24 hours, see Fig C in S1 Appendix.

### Statistics

General linear and / or mixed effect models were used to quantify the contribution of different factors to sleep timing.

Intraclass correlations (ICCs) were calculated for daily measures of mid-sleep time, sleep onset, sleep offset, sleep duration and light metrics in order to assess the within versus the between subject variance. The ICCs were calculated by first fitting a linear mixed effect model using the MATLAB fitlme function with participant as a random variable, i.e.

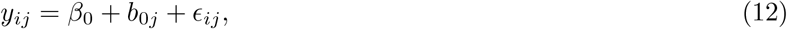

for observation *y*_*ij*_ and participant *j*, with the assumptions that *b*_0*j*_ and *ϵ*_*ij*_ are normally distributed, 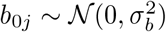 and *ϵ*_*ij*_ ∼ 𝒩 (0, *σ*^2^). Hence, 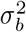 is the between participant variance and *σ*^2^ is the residual variance (within participant variance). The ICC is then given by

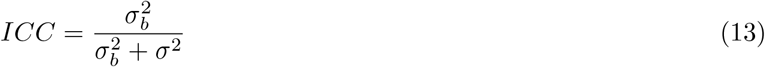

Pairwise Spearman’s rank correlations were computed using the MATLAB function corr. Significance levels were corrected for false discovery rate (FDR) using the Benjamini-Hochberg method [52] with a FDR of 5%.

## Results

### Behaviour of the HCL model: sensitivity of sleep duration and timing to model parameters and light availability patterns

Before fitting, we perform a sensitivity analysis of the nonlinear HCL model to understand the underlying mathematical structure, deduce relevant parameters and assess which parameters are identifiable from available data.

We find parameters largely separate into those that primarily affect sleep duration and those that affect sleep timing. Sleep timing is also affected by light exposure. Results are summarised in Fig 2 and Figs D and E in S1 Appendix and discussed further below.

**Fig 2.**
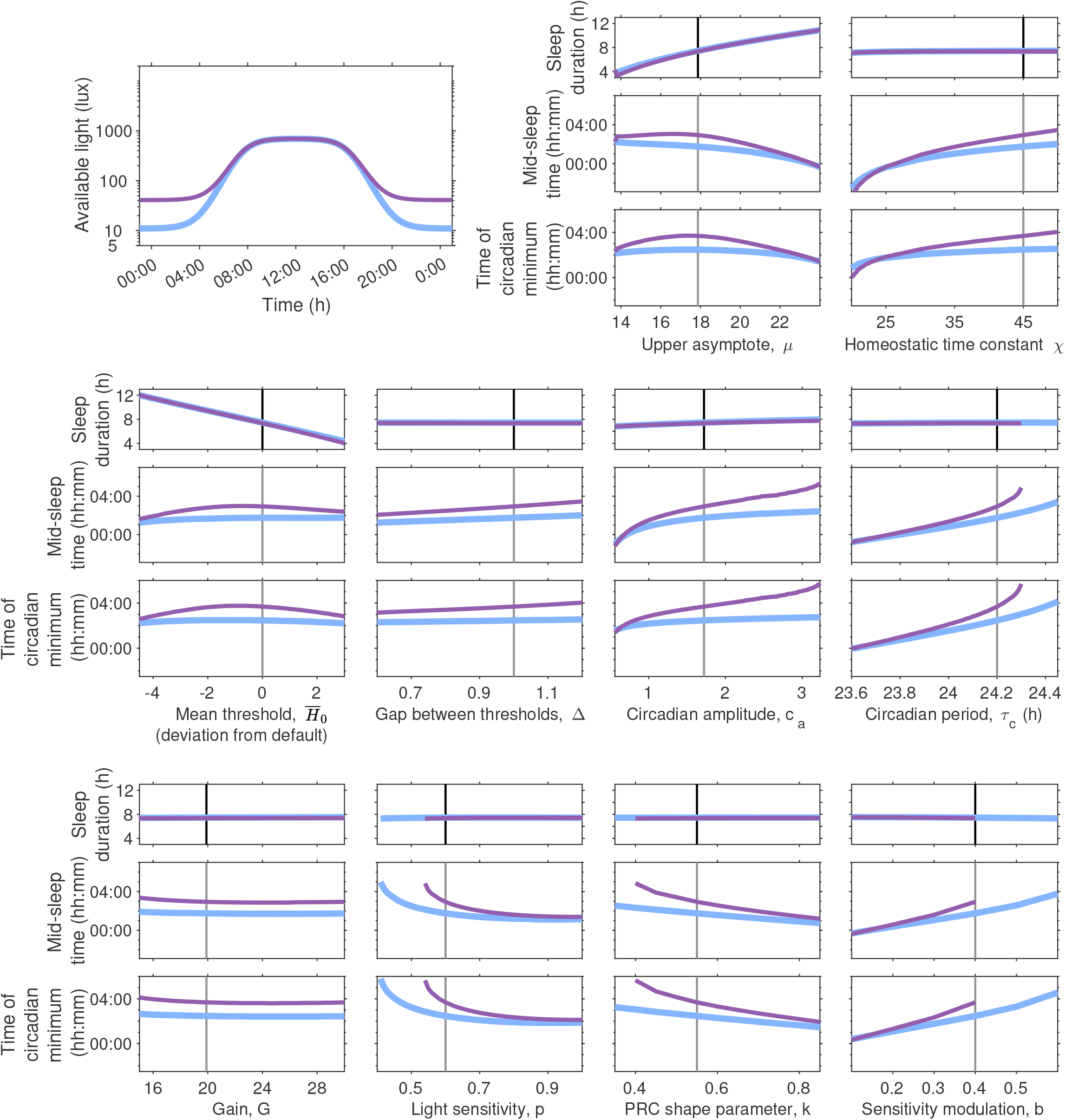
Dependence of model predicted sleep duration and timing on model parameters and light exposure. The top left panel shows two different light availability profiles. In all other groups of panels, the dependence of sleep duration (top panels), mid-sleep time (middle panels) and time of the circadian minimum (bottom panels) for the two light profiles are shown. The black vertical line indicates the default parameter values. Results are shown for all five sleep-wake parameters and five of the circadian-light parameters. The equivalent graphs for the remaining four circadian-light parameters are shown in Fig D in S1 Appendix.

#### Parameters that affect sleep duration

For the five sleep-wake regulation parameters listed in Table 1, sleep duration is primarily controlled by *μ* (the upper asymptote) and 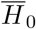 (the mean level of the wake propensity rhythm). Both of these parameters affect sleep duration through a similar mechanism. Either increasing 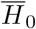 or decreasing *μ* result in a net reduction in the rate of increase of sleep homeostasis during wake. The slower rise in sleep homeostasis during wake results in longer wake periods and consequently shorter sleep periods.

The parameter *χ*, which simultaneously affects the rate of decay of sleep homeostasis during sleep and the rate of rise of sleep homeostasis during wake, has little impact on sleep duration over the range considered. Similarly, sleep duration is insensitive to the gap between the thresholds, Δ over the range considered. Circadian amplitude *c*_*a*_, has a small effect on sleep duration.

There are ten light / circadian parameters listed in Table 1, although only nine of them are independent since *f* and *τ*_*c*_ only appear in the combination *fτ*_*c*_. All of these nine parameters act to change the timing of the circadian rhythm relative to the 24-hour day with barely observable changes to the amplitude or shape of *C*(*t*) (graphs for five of these nine parameters are shown in Fig 2, the remaining four are shown in Fig D in S1 Appendix). Unless the light is sufficiently strong to change the shape of *C*(*t*), there is no mechanism for any of these parameters to change sleep duration.

#### Parameters that affect sleep timing

Each of the five sleep-wake regulation parameters affects sleep timing to a greater or lesser extent. The change in sleep timing occurs through two mechanisms. First, where there are changes to sleep duration there are consequently also changes to the duration of the light period. Second, all five parameters alter the internal relationship between sleep and circadian rhythmicity to some extent (see Fig E in S1 Appendix). The fixed light availability profile used here combined with the gating effect of sleep, mean that a shift in the circadian minimum towards the end of the sleep episode results in more bright light earlier in the circadian day. More bright light earlier in the circadian day means more light at times when light acts to speed up the circadian clock, leading to earlier sleep timing.

Each of the nine independent circadian parameters adjust sleep timing by changing the magnitude or the effect of light on the circadian pacemaker, consistent with their definitions in Table 1. The parameter *τ*_*c*_ is the intrinsic period of the van der Pol oscillator modelling the circadian pacemaker. Consistent with entrainment theory [53], sleep timing is later for longer *τ*_*c*_. For an evening light of 40 lux, the curves for the time of the circadian minimum and the mid-sleep time stop at around *τ*_*c*_ equals 24.3 h. At this point, the parameter values sit at the edge of the region of entrainment for stable periodic sleep-wake cycles. For larger values of *τ*_*c*_ the model no longer entrains. For similar reasons, the curves for *p, k, b* and *I*_0_ for an evening light level of 40 lux also stop.

The parameters *α*_0_, *p* and *I*_0_ collectively alter the magnitude of the light signal. Increasing *α*_0_ or *p* scales all values of light up. Increasing *I*_0_ scales all values of light down. However, since both the magnitude and the timing of light are important it is not necessarily obvious what the net effect of scaling light will be on sleep and circadian timing. Here, we find that for the synthetic light profiles shown in Fig 2, increasing *p* or *α*_0_ or decreasing *I*_0_ shifts sleep and circadian timing earlier.

The parameter *β* alters the rate of decay of activated photoreceptor. Larger values of *β* will in general lead to smaller values for *n* and thus larger values for 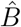 and *B. k* alters the shape of the velocity response curve that describes the effect that light has on the velocity of the clock as a function of the phase of the clock. *b* alters the timing when the clock is most sensitive to light. The parameter *γ* alters the speed at which the van der Pol oscillator returns to the limit cycle. Under the entrained conditions here with a smooth, regular periodic input it has little impact on mid-sleep timing.

#### The effect of light on sleep timing

The sensitivity results presented in Fig 2 and Fig D in S1 Appendix are shown for two different example light profiles. These two profiles have a similar shape, with light switching smoothly between a high level during daylight hours and a low level at all other times, representing 24-hour access to electric light. The duration of the daylight hours was selected to approximate the natural photoperiod around the equinoxes. We refer to the low level as ‘evening light’ because the alignment of sleep with respect to daylight hours means that the primary effect of the low level of light is after dusk. The two profiles differ in the amount of available evening light. Note that the model assumes that light is available during wake, but is turned off when the switch to sleep occurs and turned back on when the switch to wake occurs. The consequent ‘self-selection’ means that sleep timing controls the timing of light, resulting in feedback between sleep and circadian rhythmicity [13, 14]. This feedback has a ‘destabilising’ effect since even if the input light signal is periodic the signal reaching the retina will not be periodic unless sleep is periodic.

In all cases, increasing evening light shifts sleep and circadian timing later with no visible impact on sleep duration. However, the magnitude of the delay is dependent on the parameter values. For the default parameters, the increase in evening light results in a delay of approximately one hour because increasing light during the evening results in more light exposure during times when light acts to slow down the circadian clock [54]. For the intrinsic period *τ*_*c*_, longer circadian periods result in greater sensitivity to evening light, consistent with [14].

In general, the parameter sensitivity curves have steeper slopes for the higher value of evening light, meaning that sleep and circadian timing are more sensitive to the parameter values for evening light of 40 lux than of 10 lux. This is because of the gating effect of sleep on light exposure. When the evening light levels are low, the precise timing of sleep onset does not greatly alter the light exposure pattern reaching the eye and so sleep timing has little impact on circadian timing. However, when the evening light levels are high, later sleep onset leads to more light in the evening which further contributes to the delay to the circadian clock.

#### The same phenotype may have different driving factors

Fig 2 highlights that multiple model parameters lead to a particular sleep phenotype. For example, an individual may have a late sleep timing preference because of their physiology (e.g. a long intrinsic circadian period or a high sensitivity to light in the evening) or because of their pattern of light exposure. Different phenotypes and associated model parameters are summarised in Table 2.

**Table 2.** Different sleep phenotypes, associated model parameters and the physiological and environmental interpretation.

| Sleep phenotype | Physiological / environmental interpretation | Associated model parameter |
| --- | --- | --- |
| Long sleep | Rapid increase of homeostatic sleep pressure during wake.<br>High sleep need. | High value of $\mu$. |
| | Low level of wake propensity. | Low mean level of thresholds $\bar{H}_0$. |
| Short sleep | Slow increase of homeostatic sleep pressure during wake.<br>Low sleep need. | Low value of $\mu$. |
| | High level of wake propensity. | High mean level of thresholds $\bar{H}_0$. |
| Early sleep | Low circadian amplitude. | Small circadian amplitude $c_a$. |
| | Intrinsic period shorter than 24 h giving a tendency to phase advance in the absence of zeitgebers of sufficient strength. | Short $\tau_c$. |
| | Insensitive to the effect of light in the evening and / or sensitive to bright light during the day. | High value of $p$, high value of $k$. |
|  | High levels of bright light during the day and / or low levels of light in the evening. |  |
| Late sleep | High circadian amplitude. | High $c_a$. |
| | Intrinsic period longer than 24 hours giving a tendency to phase delay in the absence of zeitgebers of sufficient strength. | Long $\tau_c$. |
| | Sensitive to the effects of light in the evening and / or insensitive to the effects of bright light during the day. | Low value of $p$ or $k$ or high value of $b$. |
|  | Low levels of light during the day or too much light in the evening. |  |
| Irregular sleep | Low circadian amplitude. | Low $c_a$. |
|  | Low zeitgeber strength i.e. little variation in levels of light across the day. |  |

For given sleep timing and duration data, there may therefore be multiple ways to explain the same data. With access to individual light exposure data, it should however, be possible to separate endogenous physiological factors from exogenous environmental factors.

### Identifying endogenous factors driving sleep phenotypes: combining the HCL model with data

Next we demonstrate that we can match individual average sleep duration and timing by fitting for two pairs of parameters namely (*μ, τ*_*c*_) and (*μ, c*_*a*_). The choice of these pairs is motivated by the sensitivity analysis and aims to provide sufficient flexibility to fit to a wide range of sleep durations and mid-sleep times. Essentially, for each pair, one parameter primarily determines sleep duration (*μ*), whilst the other primarily determines sleep timing (*τ*_*c*_ or *c*_*a*_). We fit two pairs of parameters to illustrate some of the pros and cons of parameter selection.

#### Observed sleep and circadian phenotypes

In order to test the HCL model, it was important to consider individuals who exhibit a range of different phenotypes. In our cohort, sleep timing (mid-sleep ICC = 0.65, sleep onset ICC = 0.56, sleep offset ICC = 0.61), duration (ICC = 0.47) and day-to-day variability differed between individuals. Five examples are shown in Fig 3 and include: an early sleeper (mid-sleep time ± standard deviation 02:11 hh:mm ± 0:26 h:mm; sleep onset 22:55 hh:mm ± 0:30 h:mm; sleep offset 05:28 ± 0:49 h:mm); a late sleeper (mid-sleep time 04:18 hh:mm ± 0:42 h:mm; sleep onset 00:53 ± 0:47 h:mm; sleep offset 07:44 hh:mm ± 0:55 h:mm); a long sleeper (sleep duration ± standard deviation 8.92 h ± 0.92 h); a short sleeper (sleep duration 4.67 h ± 0.74 h) and an irregular sleeper (sleep duration 5.37 h ± 1.85 h). Also shown are boxplots for sleep duration and mid-sleep time for each participant and the distributions of the participant average and standard deviation in mid-sleep time and sleep duration for the whole cohort. Equivalent boxplots and distributions for sleep onset and offset are shown in Fig F in S1 Appendix.

**Fig 3.**
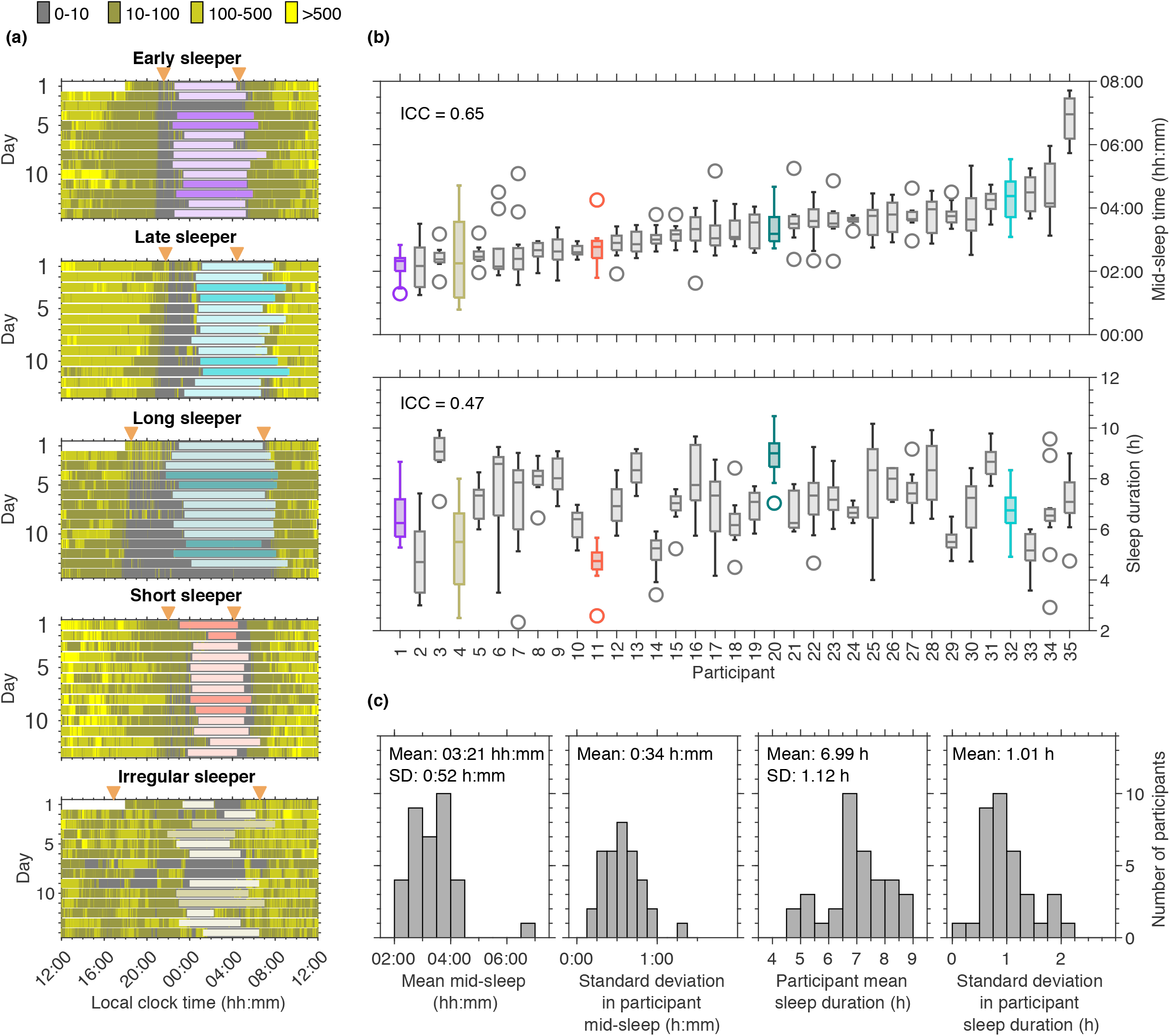
Example of field data for five different sleep phenotypes and summary information for sleep timing and duration for each participant and the cohort. Panels (a) show daily sleep timing (coloured horizontal bars, darker colouring shows sleep on Friday and Saturday nights). Light exposure is shown in the background for four different intensity bands. Dusk and dawn are indicated by the orange triangles. Panels (b) show box plots for mid-sleep timing and sleep duration respectively for each participant. Participants have been ordered according to their mean mid-sleep time. Panels (c) show the distributions of mean participant mid-sleep, standard deviation of participant mid-sleep, mean participant sleep duration, and standard deviation of participant sleep duration respectively.

#### Personalised model parameters were retrieved for each participant

Model parameters (*μ, τ*_*c*_) that matched average sleep timing and duration for every individual were found for all 34 participants. Fitting for (*μ, c*_*a*_) successfully retrieved parameters for 26 of the participants when raw light data were used and 30 of the 34 participants when imputed light data were used. The reason that not all participants were fitted when using (*μ, c*_*a*_) are discussed in the section ‘Which parameters to fit?’ below. Example fits and the corresponding personalised HCL models are shown in Fig 4. Scatter plots showing the values of all parameters can be seen in Fig 5.

**Fig 4.**
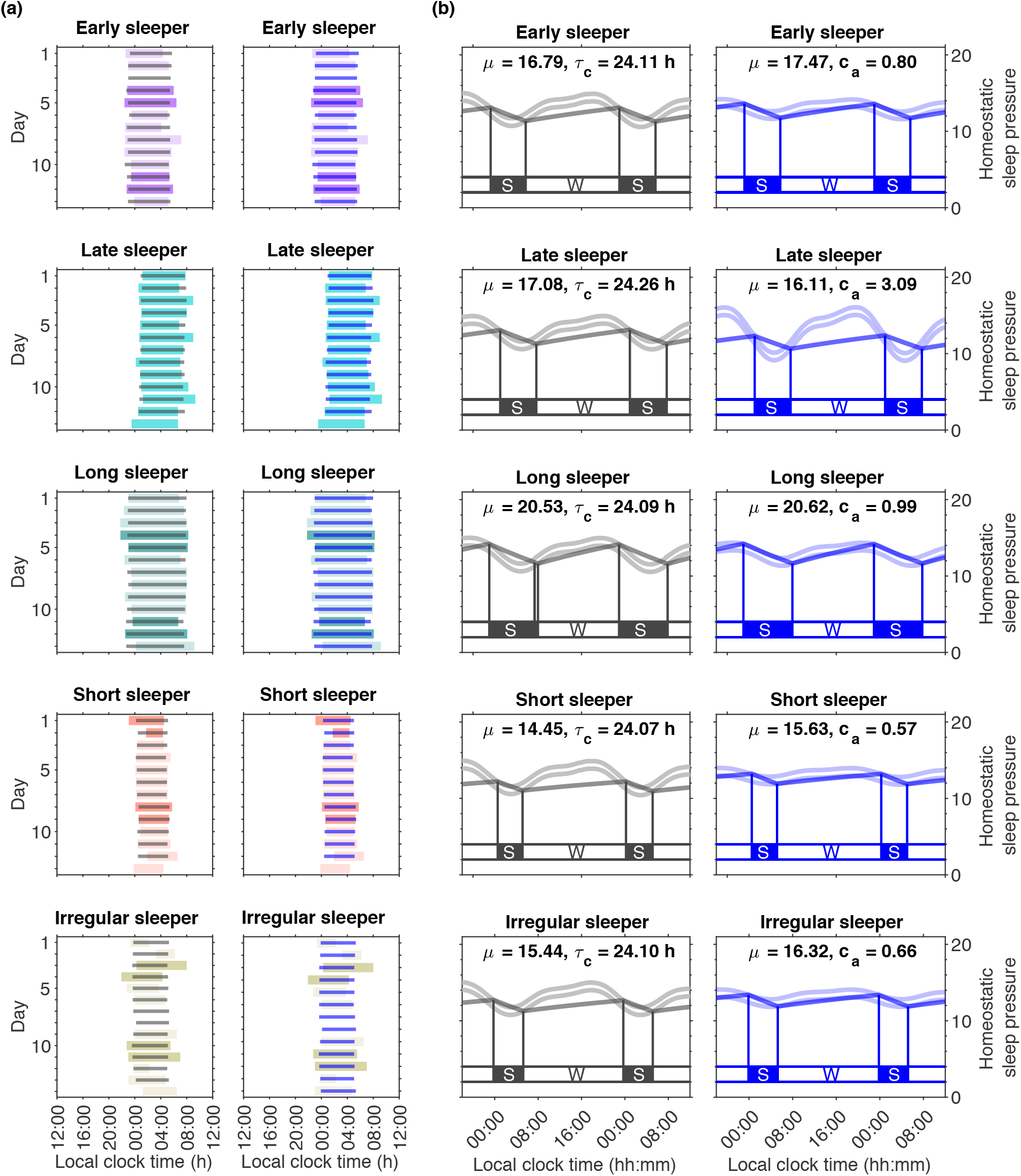
Example fits to mid-sleep and sleep duration. Panels (a) show simulated raster plots superimposed on the sleep timings (background bars coloured according to phenotype) as a result of fitting for the homeostatic parameter *μ* with the two alternative circadian timing parameters, i.e. the intrinsic circadian period, *τ*_*c*_ (grey, left hand column), and the circadian amplitude *c*_*a*_ (blue, right hand column). Panels (b) show the corresponding Homeostatic-Circadian-Light (HCL) models from fitting for (*μ, τ*_*c*_) (grey, left hand column, *c*_*a*_ fixed at the default value of 1.72) and for (*μ, c*_*a*_) (blue, right hand column, *τ*_*c*_ fixed at the default value of 24.20 h).

**Fig 5.**
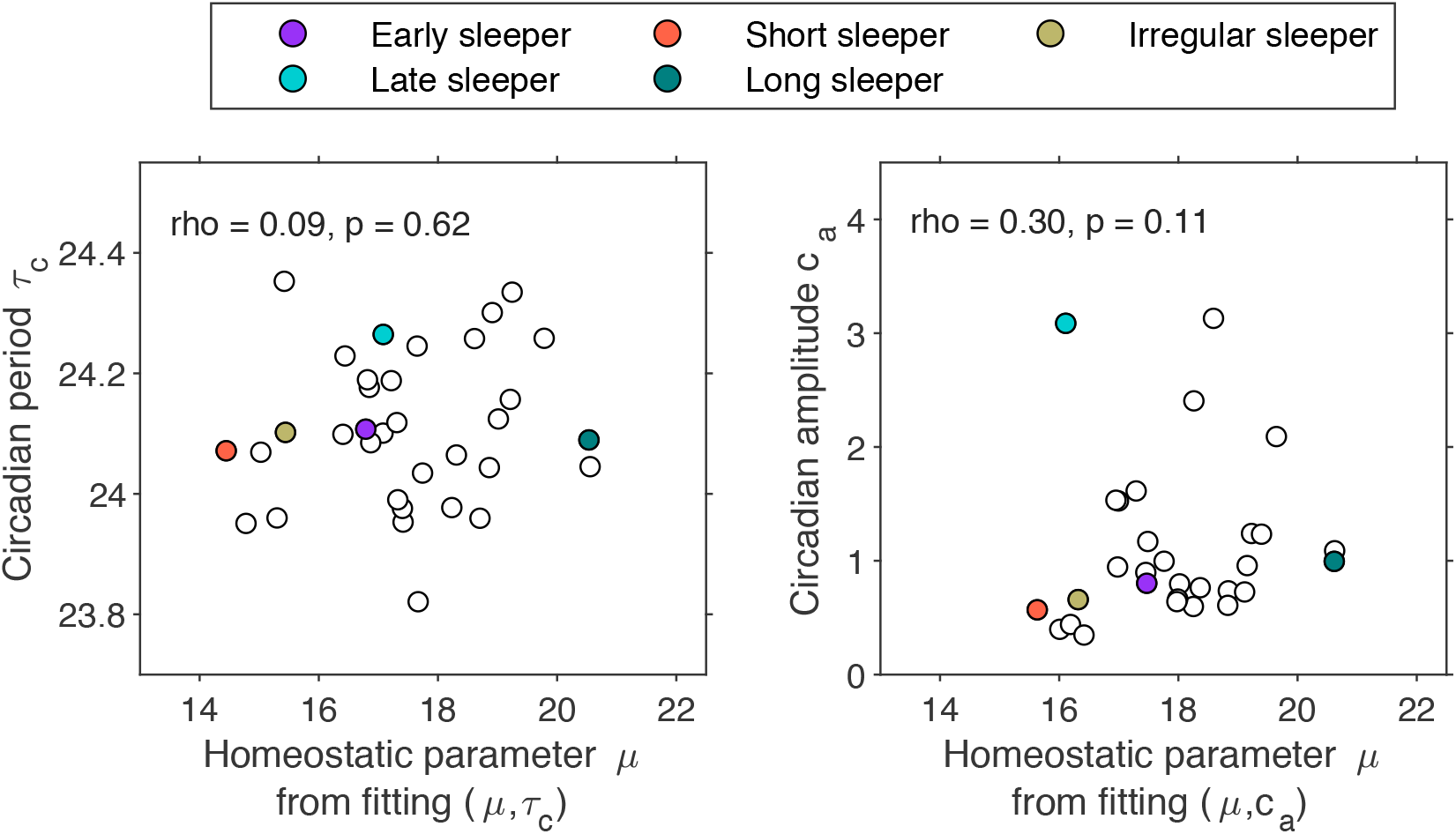
Associations between the fitted homeostatic parameter *μ* with the two alternative circadian timing parameters, i.e. the intrinsic circadian period, *τ*_*c*_ and the circadian amplitude *c*_*a*_.

For both pairs of parameters (*μ, τ*_*c*_) and (*μ, c*_*a*_), the fitted values accurately predict average sleep duration and mid-sleep timing within individuals to within the tolerances set for the optimization algorithm i.e that the residual calculated as the square root of the sum of the squares of the error in the average sleep duration and the average mid-sleep was less than two minutes, Fig G in S1 Appendix.

There is a very little association between the homeostatic parameter and the circadian period *τ*_*c*_, supporting the fact that they are capturing different phenotypic dimensions, see Fig 5. There was a weak association between the homeostatic parameter and the circadian amplitude, *c*_*a*_.

The method was robust in that fitted parameters calculated using data collected from the wrist were strongly associated with those calculated using data collected from the shoulder (Spearman’s rho for the homeostatic parameters *μ* ≥ 0.96, and those for the timing parameters of *c*_*a*_ or *τ*_*c*_ ≥ 0.79), see Figs H and I in S1 Appendix. Likewise, parameters fitted using imputed data were strongly associated with those calculated using raw data (Spearman’s rho values for the homeostatic parameters ≥ 0.97, and for the timing parameters of *τ*_*c*_ or *c*_*a*_ ≥ 0.90).

#### Relation of fitted parameters to phenotypes and physiology

Consistent with the sensitivity analysis, we find that for each pair of parameters, one parameter (*μ*) is strongly associated with sleep duration, with little association with mid-sleep time, see Table 3. Larger *μ* is associated with longer sleep duration, shifting sleep onset earlier and sleep offset later by approximately the same amount.

**Table 3.** Correlations between fitted parameters and sleep duration and timing measures and slow wave activity. Values in bold are significant (p *<* 0.05). Those with an asterisk survive FDR correction (p ≤ 0.0125).

|  | Spearman correlations (p-values) |  |  |  |  |  |  |  |  |  |
| --- | --- | --- | --- | --- | --- | --- | --- | --- | --- | --- |
|  | Mid-sleep time |  | Sleep onset |  | Sleep offset |  | Sleep duration |  | Slow wave activity |  |
| Homeostatic parameter $\mu$ ($\mu, \tau_c$ fitting) | 0.05 | (0.784) | -0.60 | (<0.001*) | 0.59 | (<0.001*) | 0.99 | (<0.001*) | 0.40 | (0.020) |
| Homeostatic parameter $\mu$ ($\mu, c_a$ fitting) | 0.01 | (0.976) | -0.67 | (<0.001*) | 0.47 | (<0.010*) | 0.95 | (<0.001*) | 0.32 | (0.083) |
| Circadian timing parameter $\tau_c$ | 0.34 | (0.053) | 0.12 | (0.497) | 0.33 | (0.055) | 0.13 | (0.464) | 0.05 | (0.780) |
| Circadian timing parameter $c_a$ | 0.27 | (0.154) | -0.22 | (0.246) | 0.52 | (0.004*) | 0.51 | (0.005*) | 0.19 | (0.320) |

Also consistent with the sensitivity analysis, *τ*_*c*_ or *c*_*a*_ are (weakly) associated with mid-sleep time, with larger values of *τ*_*c*_ and larger values of *c*_*a*_ indicative of later mid-sleep time. We note that although we consider *c*_*a*_ as a ‘timing’ parameter, as found in the sensitivity analysis, it does also affect sleep duration and indeed there is a moderate association between *c*_*a*_ and sleep duration.

The two values of the homeostatic parameters *μ* are strongly associated with each other (Spearman’s rho 0.96), as are the timing parameters *τ*_*c*_ and *c*_*a*_, (Spearman’s rho 0.84), see Fig G in S1 Appendix.

The HCL model is designed to capture the gross features of key physiological processes that regulate sleep. Since SWA activity is viewed as a marker of homeostatic sleep pressure it is interesting to note that the homeostatic parameter *μ* derived from data collected at home is positively associated with SWA derived from data collected in the laboratory, see Table 3.

The width of the distribution of model intrinsic circadian periods *τ*_*c*_ (standard deviation 0.12-0.15 h) is similar to the width of the distribution of intrinsic circadian periods found in forced desynchrony experiments (standard deviation 0.2 h, [7]), see Fig 6(a). However, since both circadian amplitude *c*_*a*_ and *τ*_*c*_ determine sleep timing, in the HCL model these interact, consequently the mean value for the distribution of fitted *τ*_*c*_ depends on the circadian amplitude. For example, when fitting for (*μ, τ*_*c*_) a default value of *c*_*a*_ = 1.72 was used. Using a larger circadian amplitude results in later sleep timing. So fixing the default circadian amplitude at a higher value will tend to shift the distribution of *τ*_*c*_ to smaller values (earlier sleep timing) to compensate. Similarly, a smaller circadian amplitude shifts the distribution of *τ*_*c*_ to higher values, see Fig 6(a). Hence, one cannot make inferences on the actual intrinsic circadian period as measured in a forced desynchrony protocol. We have kept the notation *τ*_*c*_ consistent with prior literature (e.g. [28]), but it is perhaps more appropriate to think of it as an endogenous timing parameter.

**Fig 6.**
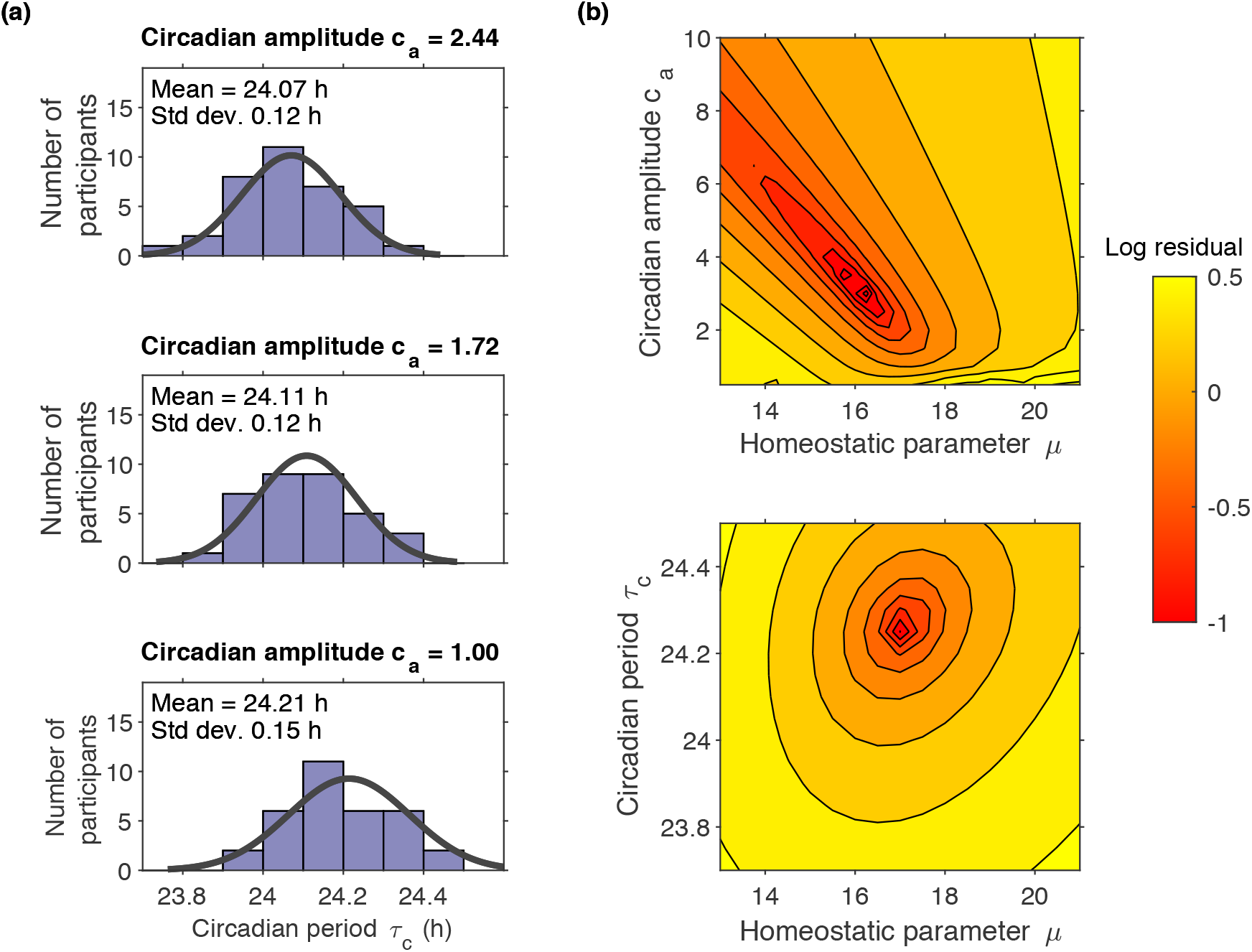
Identifiability of model parameters. Panels (a) show histograms of fitted intrinsic circadian period *τ*_*c*_ for three different choices of the circadian amplitude *c*_*a*_, highlighting the inter-dependence of these two parameters. Panels (b) show the residual for fitting simultaneously to mean sleep duration and mean mid-sleep timing for one participant as a function of (*μ, τ*_*c*_) and (*μ, c*_*a*_) i.e. the sum of the squared differences between mean sleep duration and mean simulated sleep duration and mean mid-sleep time and mean simulated mid-sleep time.

#### Day-to-day variability

Average sleep duration and mid-sleep timing are accurately matched, but Fig 4 highlights that there are differences between reported and model predicted day-to-day mid-sleep timing. Specifically, average standard deviation in participant reported mid-sleep time was 0:34 h:mm (see Fig 3), whereas model predictions were 0:08 h:mm and 0:07 h:mm for (*μ, τ*_*c*_) and (*μ, c*_*a*_) respectively. Similarly, average participant standard deviation in reported sleep duration was 1.01 h (see Fig 3), whereas model predictions were 0.10 h and 0.07 h for (*μ, τ*_*c*_) and (*μ, c*_*a*_) respectively.

Considering the day-to-day differences between predicted and reported mid-sleep time, sleep onset time, sleep offset time and sleep duration, there were no significant differences between the goodness of fit for (*μ, τ*_*c*_) and (*μ, c*_*a*_).

#### Which parameters to fit?

Overall, it was faster and more people were successfully fitted with (*μ, τ*_*c*_) than (*μ, c*_*a*_). Plotting the residual error as a function of the parameters, see Fig 6(b) for one example, illustrates the reasons. For (*μ, τ*_*c*_) there is a well-defined minimum value since sleep duration is independent of *τ*_*c*_ and mid-sleep is only weakly dependent on *μ*. For (*μ, c*_*a*_) the minimum value is much less well-defined with a line along which reducing *μ* and increasing *c*_*a*_ give similar (low) values for the residuals. The inability to fit for (*μ, c*_*a*_) for a few people is because for some light exposure patterns, the model is unable to simulate late enough sleep timing by only varying amplitude (and (*μ*)). This is because for large amplitudes, sleep timing is very insensitive to further changes in amplitude. This can be seen in the sensitivity analysis shown in Fig 2 and understood with reference to the HCL model for the late sleeper shown in Fig 6(b). In the HCL model, once the amplitude is large, at the points on the thresholds where sleep onset and offset occur, the gradient is very steep. Increasing the amplitude further makes the gradient steeper, making the time of sleep onset or sleep offset relatively insensitive to the upward or downward trajectory of the sleep homeostat.

Table 2 and the sensitivity analysis shown in Fig 2 and Fig E in S1 Appendix highlight that multiple parameters affect sleep timing. The fact that we could fit 30 of the 34 participants equally well for two pairs of parameters (*μ, c*_*a*_) and (*μ, τ*_*c*_) further highlights that it is not possible to uniquely identify a set of parameters for each participant on the basis of light, sleep timing and duration data alone. As shown in Fig 6, a slightly larger *c*_*a*_ can be compensated for by a slightly smaller value for *τ*_*c*_ (and a smaller value of *μ*). Similarly, it is not easy to distinguish between small differences in light sensitivity (e.g. the parameter *p*) and small differences in one of the other parameters that alter the effect of light on sleep timing (e.g. *c*_*a*_ and *τ*_*c*_).

On the other hand, the fact that we could not fit 4 of the 34 participants by only varying (*μ, c*_*a*_) illustrates that picking parameters with sufficient sensitivity is important. Since we were able to fit sleep timing (and duration) for all participants by varying *τ*_*c*_ (and *μ*), from this point we focus on the pair of parameters (*μ, τ*_*c*_).

### Exogenous factors determining sleep phenotypes

The endogenous parameters found by fitting the HCL model were strongly associated to sleep duration, sleep onset and sleep offset, but the association with mid-sleep timing was weak. Given the acknowledged importance of light in determining the timing of the biological clock and sleep, it is interesting to examine if individual patterns of light exposure correlate with sleep phenotypes.

#### Observed light exposure patterns

Light exposure patterns varied between individuals, see Fig 7, where light exposure patterns for the five example sleep and circadian phenotypes are shown. Also shown are boxplots indicating the daily hours of bright light and daily measure of the effect of light on the biological clock for each participant and distributions of the participant average and standard deviation in both light measures. An average of less than one hour of bright light (light *>* 500 lux) a day was recorded at the wrist for most people (24 individuals i.e. 71%). Equivalent boxplots and distributions for the remaining four light metrics which further quantify overall light exposure and timing of light exposure are shown in the Figs J and K in S1 Appendix.

**Fig 7.**
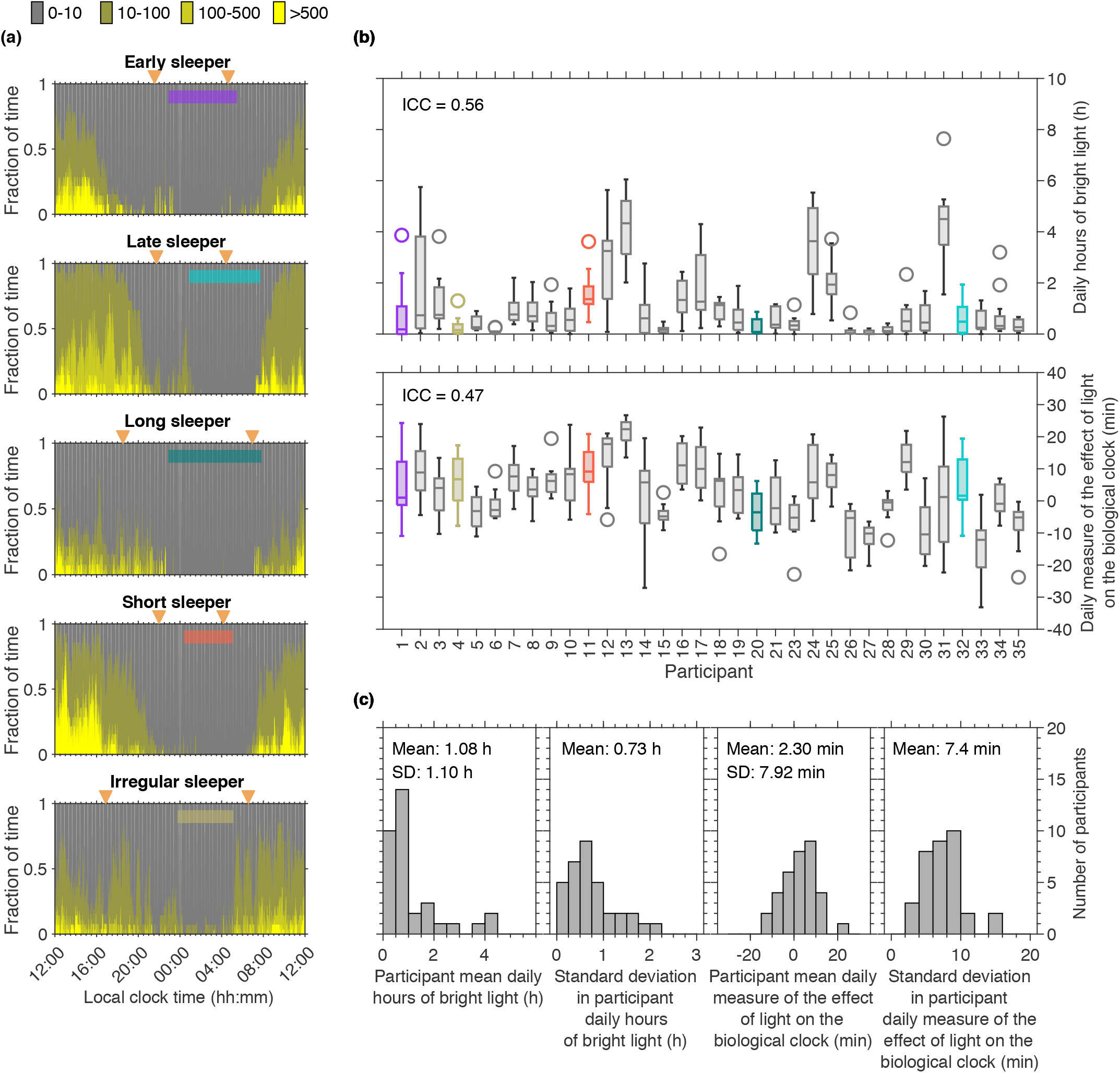
Example light data for five different sleep phenotypes and summary information for each participant and the cohort. Panels (a) show examples of light exposure patterns for five different participants. The regions show the fraction of time spent in each of four intensity bands. Average sleep timing is marked by the coloured horizontal bars. Dusk and dawn are indicated by the orange triangles. Panels (b) show box plots for the daily hours of bright light and the daily measure of the effect of light on the circadian clock respectively for each participant. Participants are ordered by their mean mid-sleep time, as in Fig 3. Panels (c) show the distributions of the mean and standard deviation of participant daily hours of bright light and the mean and standard deviation of the participant daily measure of the effect of light on the biological clock.

#### Associations between sleep timing and duration and light exposure

Mid-sleep time was not associated with any of the three metrics which quantify the amount of light received, i.e. the mean lux value, the geometric mean value or the mean number of daily hours of bright light (light *>* 500 lux) (Spearman’s rho all of magnitude less than or equal to 0.17), see Table B in S1 Appendix. Mid-sleep was moderately associated with the two measures of the timing of light i.e. the time of day at which half the total daily amount of light (lux) has been received (Spearman’s rho 0.47) and the time of day at which half the total daily amount of light (log (lux+1) has been received (Spearman’s rho 0.41). The associations are such that later light exposure is associated with later mid-sleep time. The novel measure of the biological effect of light was also moderately assciated with mid-sleep time (Spearman’s rho -0.41).

All measures that quantify the amount of light are significantly associated with each other (p*<*0.001), as are the measures that quantify the timing of light (p*<*0.001), see Table B in S1 Appendix. The novel measure of the biological effect of light was found to be moderate to strongly associated with all measures of the amount of light (Spearman’s rho from 0.53 to 0.71) and with one of the timing measures (time of half-light measured in lux, Spearman’s rho -0.68).

Sleep duration was not found to associate with any of the six light metrics (Spearman’s rho all of magnitude less than or equal to 0.19), see Table B in S1 Appendix.

### Relative contribution of endogenous physiological and exogenous environmental drivers of sleep phenotypes

To summarise, individual differences in sleep timing and duration were successfully fitted by the HCL model. Individual differences in sleep duration were found to be primarily driven by the modelled homeostatic sleep process (see Table 3). However, for individual differences in mid-sleep timing the explanation is more complex and is a result of the interaction of physiological factors and light exposure patterns. In a first approach to determine at an individual level the relative contribution of physiological factors and light exposure patterns we have found model parameters describing endogenous physiological factors and used a variety of measures to quantify exogenous light exposure.

Physiological factors driving sleep timing are captured by the fitted model circadian timing parameter, *τ*_*c*_.

However, interestingly, we found that *τ*_*c*_ is only weakly associated with mid-sleep timing, only explaining a small amount of the variance, see Table 3 and Fig 8(a). For example, although the late sleeper labelled as ‘late phenotype (physiology)’ in Fig 8(a) has a greater value of *τ*_*c*_ than the ‘early phenotype’, there are those with a relatively small value of *τ*_*c*_ who have a similarly late mid-sleep time, for example the individual labelled as ‘late phenotype (environmental light)’.

**Fig 8.**
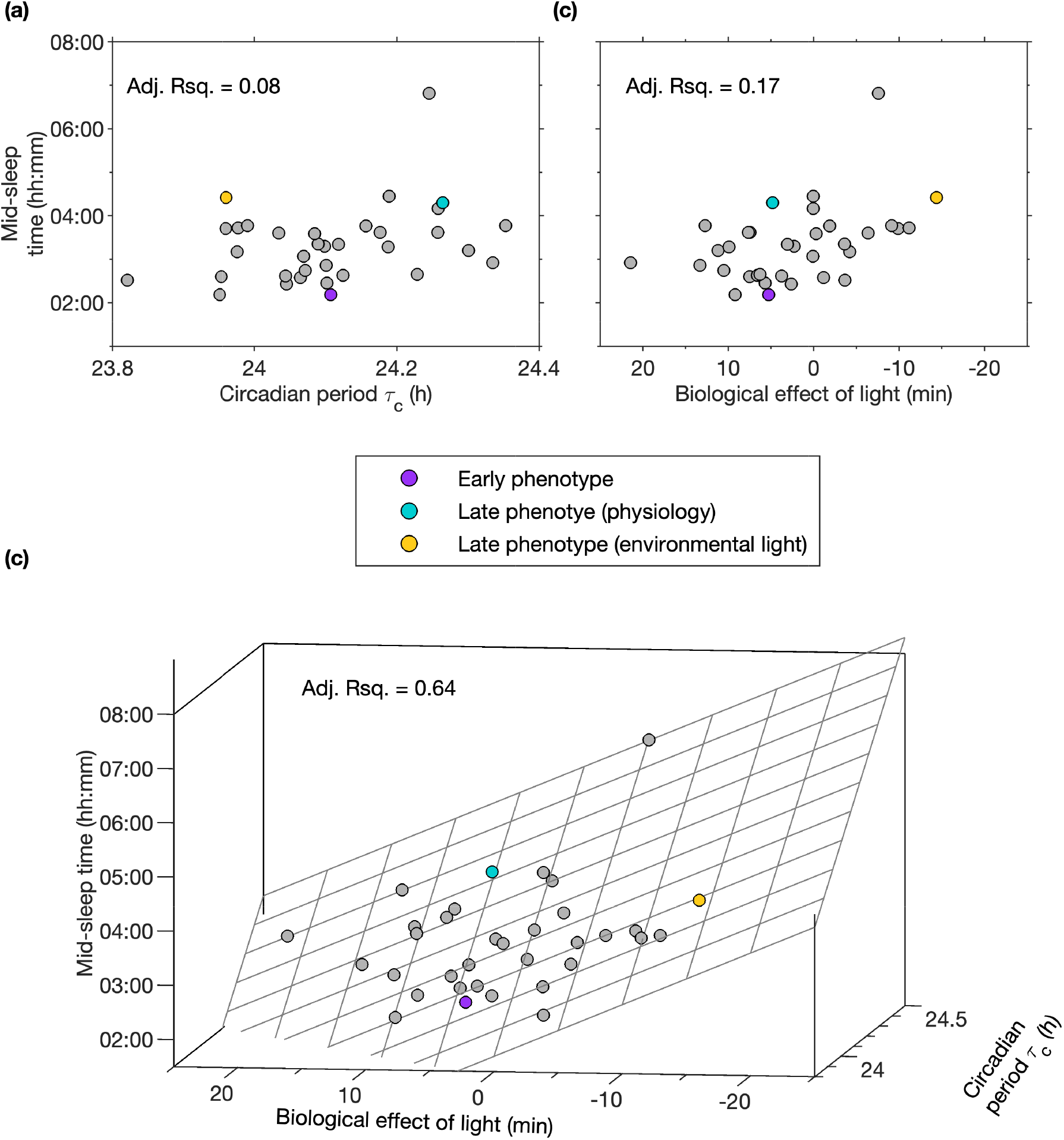
Quantifying the effect of environment versus physiology in the determination of mid-sleep time. Panel (a) shows that the fitted physiologically-motivated timing parameter, the intrinsic circadian period *τ*_*c*_, also only explains a small amount of the between participant variance. Similarly, panel (b) shows that considering the novel metric of the biological effect of light also explains only a small amount of the between participant mid-sleep time. In contrast, combining environmental and physiological variables (panel (c)), explains a large amount of the variance.

Similarly, in spite of the acknowledged importance of light as the primary zeitgeber for the human biological clock, mid-sleep timing was not associated with measures of total environmental light exposure and was only moderately associated with measures of the timing of light (see Table B in S1 Appendix). As shown in Fig 8(b), our novel metric of the biological effect of light also only explains a small amount of the variance in mid-sleep time, with the early phenotype and late phenotype (physiology) examples having similar values for the biological effect of light.

However, taken together, the fitted parameter *τ*_*c*_ and our novel metric of light timing explain a large amount of the variance (adjusted Rsquared, 0.64), see Fig 8(c). Here we see that our modelling approach suggests that of the two highlighted late sleep examples, one has late phenotype as a result of their physiology and one has a late phenotype as a result of their light exposure pattern.

Shown in Fig 8(c) are the mid-sleep timing data along with the fitted general linear model,

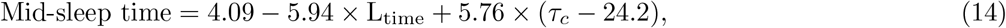

where *L*_time_ is the measure of the biological effect of light in hours. This model estimates the deviation from a mid-sleep time of 4.09 h that occurs either as a result of a deviation of the fitted intrinsic period from 24.2 hours or as a result of an environmental light signal that for the average person would advance / delay the biological clock relative to a 24-hour day. Including higher order interaction terms does not improve the fit.

The novel light timing metric combined with *τ*_*c*_ explains more of the variance than combining either of the other two light timing metrics with *τ*_*c*_. (Fitting for *τ*_*c*_ with the time of half light gives an adjusted rsquared value of 0.49, and with the time of half-loglight gives an adjusted rsquared value of 0.35.)

### Light intervention and phenotype: an example

Here, we illustrate that the underlying factors determining a given phenotype may have implications for the effects of light interventions. The predicted effect of two typical types of light intervention is shown in Fig 9 for the participants that were classified as late phenotype (environmental light) and late phenotype (physiology). One intervention is evening light ‘reduction’, in which all values of the available light between 18:00 and 06:00 were set to 4 lux (the model turns light off during predicted sleep). The other is morning light ‘enhancement’ in which all values of the available light between 9:30 and 10:00 were set to 1000 lux. All other values were left at their original recorded values. The simulations suggest that both light interventions will have a bigger impact on the indivdual for whom it was suggested that the environmental light was the main contributor to their late phenotype.

**Fig 9.**
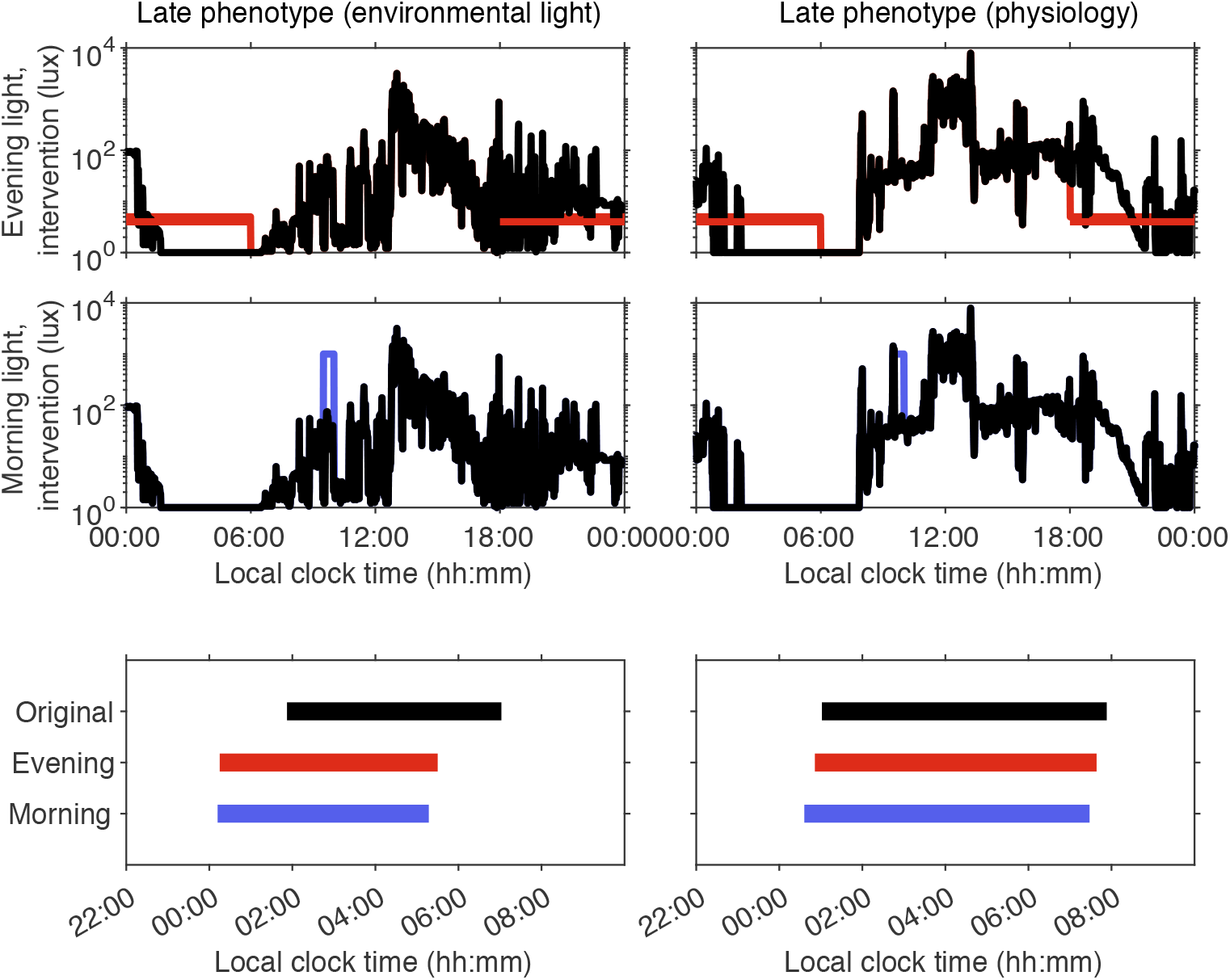
Effect of light interventions. The upper panels show one day of the original time series of the recorded light. The available light for the evening and the morning light intervention are shown in red / blue respectively. The bottom panels show the predicted sleep timing for each of the three cases (black, default; red, evening light reduction; blue, morning light enhancement). Predicted sleep timing is the average over a two week period starting two weeks after the beginning of the intervention.

## Discussion

Individuals may have preferred sleep timing that conflicts with their work schedules or personal desired sleep timing. But preferred sleep timing is a result of both endogenous physiological factors and exogenous light exposure patterns. In order to design personalised interventions to either advance or delay sleep timing an important step is to be able to dissect the relative contributions of these separate drivers to sleep phenotype. While changing endogenous physiology drivers is challenging, exogenous light exposure patterns can readily be altered and offer a practical solution for implementation of targeted interventions at scale.

Here we have developed a new simplified model for the Homeostatic-Circadian-Light (HCL) regulation of sleep timing and duration. Critically, we have demonstrated that combining the model with 7-10 days of measured light exposure patterns we can accurately find personalised parameters that match individual mean sleep-timing and duration. Parameter values are robust in that they are not strongly dependent on the position of the light sensor (wrist or shoulder) or whether raw or imputed light data are used. In addition, motivated by the fact that current widely used metrics do not capture both the intensity and time-of-day effect of light on the timing of the biological clock, we constructed a novel light metric based on the phase response to light as encoded in a van der Pol oscillator model of the circadian pacemaker [27]. The identification of personalised model parameters along with the use of the novel measure of the effect of light on the biological clock enabled us to suggest, for the first time, the relative contributions of endogenous physiological versus exogenous environmental light drivers for mid-sleep timing.

The HCL model could readily be combined with wearables to provide individual advice on appropriate light availability patterns to achieve a target preferred sleep timing, as envisaged in the interventional framework proposed in [30].

### Relation of the HCL model to other models

The HCL model was motivated by elements of the original two-process model [23] along with elements of recent neuronal models [14, 28]. Critically, unlike the two-process model it includes light to allow for the modelling of sleep-timing relative to the 24-hour day. Unlike Kronauer-type models, the HCL models both circadian rhythmicity and sleep. Compared with recent neuronal models, the HCL model is simpler (four instead of six first order differential equations, 14 instead of 23 parameters), and computationally faster. For consistency with our previous work, we checked that the HCL model gave similar results to our work on neuronal models, for example reproducing similar tongue-like entrainment regions as computed in [14] (see Fig L in S1 Appendix) and similar results to those reported in [30] when fitting to data from people living with schizophrenia and healthy unemployed controls. We found integrating the HCL model to be typically twice as fast as integrating our neuronal model.

The relative simplicity of the HCL model makes it straightforward to understand the role of each parameter, see Fig 2 and Table 2. In addition the form of the coupling between sleep and circadian rhythmicity has been updated to better reflect data on the circadian variation of wakefulness, [6]. The model can also easily be extended to include the role of social constraints (see section M and Fig M in S1 Appendix).

Nine of the parameter relate to light-circadian aspect of the model. In spite of the number of parameters, we elected to retain the circadian-light model as specified by Kronauer and colleagues [26, 27]. A recent systematic study suggests that simplifying the circadian-light model results in a poorer fit to circadian phase data [55]. Another popular version of the Kronauer model is that produced by St Hilaire and co-workers [56]. Given the similarities between the different models, we would expect results to be similar whichever version is used. Although we note that if a different light-circadian model was used then the form of the sleep-circadian coupling function *C*(*t*) given in equation (11) would also need to be updated.

### Identifying factors underlying sleep phenotypes: challenges

Based on our sensitivity analysis, we fitted for one parameter encoding sleep duration (e.g. *μ*) and one encoding sleep timing (e.g. *c*_*a*_ or *τ*_*c*_). Over several weeks, with sufficiently variable light from one week to the next, it may be possible to capture the nuanced different dynamic responses that may occur with different parameter combinations. Further types of data could enable the identification of more parameters. For example, knowledge of individual alignment of circadian phase with sleep could enable fitting of one additional sleep-wake parameter. Day-to-day variation in circadian phase could enable both a light sensitivity parameter and intrinsic circadian period to be determined as suggested for light-circadian models in [57].

### Limitations

The current work is based on only one to two weeks of data. As longer timeseries become available, the same approaches can be used to assess whether model parameters vary with time. The current HCL model, like the neuronal models, captures individual differences in the average timing and duration of sleep but not the night-to-night variation in sleep timing and duration. The reason that current deterministic sleep models do not capture night-to-night variation in sleep timing is that, in the models, the only difference between one day and the next is the difference in the light exposure pattern. Different light exposure patterns on different days introduces a small day-to-day variation in the timing of the circadian rhythm, which is translated into a day-to-day variation in sleep timing. In order for the HCL model (or any of the neuronal models) to predict greater night-to-night variation in sleep timing there are then a number of different options. For example, night-to-night variation could be modelled by the inclusion of stochasticity to capture momentary changes in alertness (e.g. see [58]) or further knowledge on the reasons that determine when people go to bed. Including circadian amplitude dependence on light [59] may also be important and enable reported dependence of sleep duration on light [60] to be captured. Finally, the current work uses lux as the measure for the effects of light on the circadian clock. The discovery of the contribution of the photopigment melanopsin to the regulation of circadian rhythmicity implies that melanopic lux rather than photopic lux may be the relevant measure [61], although we do not expect this to substantially affect our results.

### Outlook

Our modelling approach suggests that it is possible to separate endogenous physiological drivers from exogenous environmental drivers for different sleep phenotypes. This implies that there is considerable scope for appropriate models to be incorporated in clinical guidance and intervention design for sleep disorders. Models could even be implemented as standard in smart watch software for general guidance for day-to-day living or coupled to human centric lighting systems to automatically tune the light environment to optimise the alignment of sleep and circadian rhythmicity to a desired time.

## Acknowledgements

The authors would like to thank the study participants in the sleep study and all others responsible for collecting and collating the data including Dr Hana Hassanin, Damion Lambert, Surrey Clinical Research Facility staff, and Daniel Barrett. We thank Giuseppe Atzori for scoring the PSG recordings.

This work is part of the sleep and circadian disturbance project (PI DJD) supported by the UK Dementia Research Institute which receives its funding from DRI Ltd, funded by the UK Medical Research Council, Alzheimer’s Society and Alzheimer’s Research UK. The funders had no role in study design, data collection and Analysis, decision to publish or preparation of the manuscript.

## Author contributions

Conceptualization (ACS, DJD); data curation (TRG, KKGR, VLR, CdM); formal analysis (ACS, TRG, SFC); funding acquisition (DJD); investigation (ACS, TRG, SFC, VLR, KKGR, CdM); methodology (ACS, DJD); software (ACS, TRG, SFC); visualization (ACS, DJD, TRG); writing – original draft preparation (ACS, DJD); writing – review & editing (ACS, DJD, TRG, SFC, KKGR, VLR, CdM).

## Data availability statement

Software and data required for validation and replication of the results in this paper are available here: https://github.com/anneskeldon/Homeostatic_circadian_light_model-factors_driving_sleep_phenotypes

## S1 Appendix caption

The S1 Appendix includes:

**Section A** Association of the Homeostatic-Circadian-Light model with neuronal models

**Section B** Derivation of the threshold

**Section C** Comparison of the parameter regime of the HCL model with that of the original two process model

**Section D** Light metrics

**Section E** Parameter sensitivity for light parameters

**Section F** Phase angle between the time of mid-sleep and the circadian minimum

**Section G** Sleep onset and offset

**Section H** Accuracy and correlation of fitted parameters

**Section I** Sensitivity of the fitted parameters to position of the light sensor and to data imputation

**Section J** Amount and timing of light

**Section K** Associations between collected data on sleep timing and duration and timing of light

**Section L** Entrainment tongues.

**Section M** Modelling of social constraints.

## Figure legends for figures contained in S1 Appendix

**Fig A Derivation of the circadian drive for wakefulness by fitting to data**. In the top left panel data for the circadian rhythm of wakefulness are reproduced from Dijk et al, J Physiol, 516:611-627, 1999. Data for each group of participants were z-scored to remove the difference in mean level (central left panel). The bottom left panel shows the output variables *x* and *y* from the van der Pol oscillator. The right hand panels show the results of fitting to linear, quadratic and cubic functions of *x* and *y*.

**Fig B Comparison of the parameters for the circadian amplitude and homeostatic time constants for the original two-process model with the Homeostatic-Circadian-Light (HCL) model**. Left hand top panel shows a simulation using typical values for the original two-process model as compared with typical values for the HCL model. For ease of comparison, the HCL model has been scaled in the same way as the two-process model so that the upper asymptote is one. In addition, since the two-parameter model does not include a circadian-light interaction, the thresholds are here represented by sinusoids. In order to further highlight the differences in the homeostatic parameters, simulations for a zero circadian amplitude are shown in the right hand panels for the two different parameter regimes. The natural period of the sleep-wake cycle *T*_nat_ is much shorter for the HCL model.

**Fig C Light metrics**. (a) Lux levels for one day for one participant plotted on a log scale in the top panel and log (lux+1) in the bottom panel. The horizontal blue line marks the average lux value. The horizontal red line marks the average log (lux+1) value. If the average log (lux+1) is *m*, then the geometric mean is given by 10^*m*^. The vertical dashed lines mark the time of half light and time of half log light respectively. As shown in panels (b), these are calculated as the time at which half the daily cumulative value is reached for each day. Here, we have taken the day to run from midnight to midnight, but for populations who typically go to bed substantially after midnight calculating the cumulative time from, for example 04:00, may be more appropriate. (c) When light is fed into the forced van der Pol oscillator, it affects the time course of the van der Pol oscillator variables *x* and *y*. After integrating for 24 hours, the circadian oscillator will have either completed exactly one cycle, less than one cycle (as shown in the far right panel of (c)) or more than one cycle. In this example, in 24 hours, the circadian clock has only completed 98.13% of a cycle, so is 27 minutes slow. In our new light metric this is recorded as -27.

**Fig D Dependence of model predicted sleep duration and timing on model parameters and light exposure**. In each group of panels, the dependence of sleep duration (top panels), mid-sleep time (middle panels) and time of the circadian minimum (bottom panels) for the two light availability profiles given in Fig 2 are shown. The black vertical line indicates the default parameter values. The equivalent graphs for the five sleep-wake parameters and the remaining five circadian-light parameters are shown in Fig 2.

**Fig E Phase angle between mid-sleep and the circadian minimum**. Each panel shows the number of hours from mid-sleep to the circadian minimum for the five sleep-wake parameters and the circadian period *τ*_*c*_ for the two light availability profiles given in Fig 2. The black vertical line indicates the default parameter values. Like *τ*_*c*_, none of the other circadian-light parameters alter the phase relationship between mid-sleep and the circadian minimum so they are not shown.

**Fig F Summary information for sleep onset and offset for each participant and the cohort**. Top and centre panels show box plots for sleep offset and onset respectively for each participant. Bottom panels show the distributions of mean participant sleep offset, standard deviation of participant sleep offset, mean participant sleep onset and standard deviation of participant sleep onset.

**Fig G Accuracy and correlation of fitted parameters**. Fitted parameters accurately predict participant average sleep duration and mid-sleep timing, as shown by the very high correlation between observed and fitted values and the size of the residuals, shown in panels (a) for the pair of parameters (*μ, τ*_*c*_) are fitted and in (b) for the pair of parameters (*μ, c*_*a*_). The residuals were calculated using equation (S17).

**Fig H Sensitivity of fitted parameters** (*μ, τ*_*c*_) **to position of the light sensor and imputation of light data**. The top panel shows two representative days of light exposure data, collected simultaneously from the wrist and the shoulder from one participant. The central two panels show the correlation between parameters found by fitting using light collected from the wrist versus light collected from the shoulder. The lower two panels show the correlation between parameters found by fitting using raw light data versus those using imputed light data. Here, all correlations are a result of fitting for the pair of parameters (*μ, τ*_*c*_). The equivalent correlations for the pair of parameters (*μ, c*_*a*_) are shown in Fig I.

**Fig I Sensitivity of fitted parameters** (*μ, c*_*a*_) **to position of the light sensor and imputation of light data**. The top two panels show the correlation between parameters found by fitting using light collected from the wrist versus light collected from the shoulder. The lower two panels show the correlation between parameters found by fitting using raw light data versus those using imputed light data. Here, all correlations are a result of fitting for the pair of parameters (*μ, c*_*a*_).

**Fig J Summary information for measures of the amount of light exposure for each participant and the cohort**. Panels (a) show box plots for the daily mean light exposure level and the daily geometric mean of light exposure respectively for each participant. Panels (b) show the distributions of the mean and standard deviation of participant mean daily light and the mean and standard deviation of the participant geometric mean daily light.

**Fig K Summary information for measures of the timing of light exposure for each participant and the cohort**. Panels (a) show box plots for the time at which half the total daily light exposure was received in lux and log (lux +1) respectively for each participant. Panels (b) show the distributions of the mean and standard deviation of participant mean time of half light (lux) and the mean and standard deviation of the participant mean time of half light (log (lux+1)).

**Fig L Entrainment tongues** for the neuronal model in Skeldon et al, Sci. Rep. 7:45158, 2017 (left panel) and the HCL model (right panel). In both cases, the same input light profile was used – the same as the form of the light profile used for the sensitivity analysis and shown in Fig. 3 in the main text. The shading indicates mid-sleep time. The ‘evening light’ was set to 40. This means that when the ‘day light’ is also 40, the external forcing frequency takes a fixed value of 40 throughout the day and night and entrainment does not occur. For values of day light less than 40, light is brighter at night during the night than during the day and entrainment occurs to a pattern in which the sleep period is during the day (the yellow / red shaded regions).

**Fig M Modelling of social constraints and the concept of ‘wake effort’**.

## Table legends for tables contained in S1 Appendix

**Table A Threshold parameter values**

**Table B Associations between collected data on sleep timing and duration and timing of light**. The lower triangle contains the correlation coefficients (Spearman’s rho). The upper triangle indicates the p-value, shaded according to whether or not they are significant under false discovery rate correction (p *<* 0.0011). Values in grey are not significant. Associations are shown for the N=34 participants in whom light data were collected from the wrist.

