## Supplementary Material for "Method to determine whether sleep phenotypes are driven by endogenous circadian rhythms or environmental light by combining longitudinal data and personalised mathematical models"

### Contents

**A Association of the Homeostatic-Circadian-Light model with neuronal models** **7**

A.1 Hard switch approximation to the firing rate function and equivalence to the HCL model . 7

A.2 Soft switch slow manifold approximation and equivalence to the HCL model . . . . . . . . 9

**B Derivation of threshold** **9**

**C Comparison of the parameter regime of the HCL model with that of the original two process model** **10**

**D Light metrics** **13**

**E Parameter sensitivity for light parameters** **14**

**F Phase angle between the time of mid-sleep and the circadian minimum** **14**

**G Sleep onset and offset** **15**

**H Accuracy and correlation of fitted parameters** **16**

**I Sensitivity of the fitted parameters to position of the light sensor and to data imputation** **18**

**J Amount and timing of light** **21**

**K Associations between collected data on sleep timing and duration and timing of light** **23**

**L Entrainment tongues** **24**

**M Modelling of social constraints** **24**

### List of Figures

A **Derivation of the circadian drive for wakefulness by fitting to data.** In the top left panel data for the circadian rhythm of wakefulness are reproduced from [3]. Data for each group of participants were z-scored to remove the difference in mean level (central left panel). The bottom left panel shows the output variables $x$ and $y$ from the van der Pol oscillator. The right hand panels show the results of fitting to linear, quadratic and cubic functions of $x$ and $y$. . . . . . . . . . . . . . . . . . . . . . . . . . . . . . . . . . . . . . . . . . . . 9

### A Association of the Homeostatic-Circadian-Light model with neuronal models

Here, following the argument in [1], we discuss the relation between the Homeostatic-Circadian-Light model with the neuronal model of Phillips and Robinson [2].

The neuronal model of Phillips and Robinson considers two mutually inhibitory populations of neurons modelling sleep promoting neurons, primarily located in the ventrolateral preoptic and wake promoting nuerons such as monoaminergic neurons. The populations of neurons are represented by their mean cell body potential relative to rest, $V_j$ for $j = s, w$, where $s$ represents the sleep promoting neurons and $w$ represents the wake promoting neurons.

The neuronal dynamics are represented by

$$\tau \dot{V}_s + V_s = -\nu_{sw} Q_w + D_s, \tag{S1}$$

$$\tau \dot{V}_w + V_w = -\nu_{ws} Q_s + D_w, \tag{S2}$$

where $Q_j(V_j)$ are switch-like functions describing how the firing rates of the neuronal populations depend on the potentials $V_j$. The terms $D_s(t)$ and $D_w$ represent 'drives' to sleep and wake-promoting neurons, respectively, where it is assumed that $D_w$ is constant while $D_s(t)$ is given by

$$D_s = \nu_{sh} H - \nu_{sc} C(t) - A_s,$$

Here, $H(t)$ is the homeostatic sleep pressure, $C(t)$ represents the circadian drive for wakefulness. and $A_s$ is assumed to be constant.

In the original Phillips-Robinson model, $Q_j$ was taken to be the soft-switch

$$Q_j = \frac{Q_{\max}}{1 + \exp[-(V_j - \theta)/\sigma']}, \tag{S3}$$

where $Q_{\max}$ is the maximum firing rate and $\theta$ is the mean firing threshold relative to resting. The function $Q_j$ is a sigmoid function, which is close to zero for all negative values of $V_j$ and then saturates exponentially fast to $Q_{\max}$. The homeostatic component of the drive, $H$ is modelled by

$$\chi \dot{H} + H = \bar{\mu} Q_w, \tag{S4}$$

Sleep was defined as occurring when the firing rate of the wake promoting neurons dropped below a critical threshold defined as $\theta = 1$.

#### A.1 Hard switch approximation to the firing rate function and equivalence to the HCL model

An alternative hard switch approximation for the firing functions $Q_j(V_j)$ was used in [1],

$$Q_j = \tilde{Q}_{\max} \mathcal{H}(V_j - \hat{\theta}_S). \tag{S5}$$

The equation for homeostatic sleep pressure then becomes

$$\chi \dot{H} + H = \bar{\mu}\tilde{Q}_{\max}\mathcal{H}(V_w - \hat{\theta}_S). \tag{S6}$$

Here, the assumption is that the switch from wake to sleep is instantaneous, so that during wake $V_w > \hat{\theta}_S$ and $V_s < \hat{\theta}_S$, while during sleep $V_w < \hat{\theta}_S$ and $V_s > \hat{\theta}_S$. Switching from sleep to wake and vice versa occurs when $V_s = \hat{\theta}_S$.

Now, the typical timescales for the neuronal dynamics, given by $\tau$ are much faster than the typical timescales for the sleep homeostatic process, given by $\chi$. Hence, following [1], a small parameter $\epsilon = \tau/\chi$, the fast time $\hat{t} = t/\epsilon$ and the slow time $T = t$, $d/d\hat{t} = \epsilon\, d/dt$ and $d/dT = /dt$ are introduced. At $O(1)$ (slow time) equations (S1), (S2) and (S4) become

$$\begin{aligned} V_s &= -\nu_{sw}\tilde{Q}_{\max}\mathcal{H}(V_w - \hat{\theta}_S) + D_s(T), \\ V_w &= -\nu_{ws}\tilde{Q}_{\max}\mathcal{H}(V_s - \hat{\theta}_S) + D_w, \\ \chi \dot{H} + H &= \bar{\mu}\tilde{Q}_{\max}\mathcal{H}(V_w - \hat{\theta}_S), \end{aligned} \tag{S7}$$

where

$$D_w = \nu_{sh}H - \nu_{sc}C(T) - A_s.$$

Hence, during wake,

$$V_s = -\nu_{sw}\tilde{Q}_{\max} + \nu_{sh}H - \nu_{sc}C(T) - A_s, \tag{S8}$$

$$V_w = D_w, \tag{S9}$$

$$H = \bar{\mu}\tilde{Q}_{\max} + \left(H(t_{\text{off}}) - \bar{\mu}\tilde{Q}_{\max}\right) \mathrm{e}^{(T_{\text{off}} - T)/\chi}. \tag{S10}$$

where $T_{\text{off}}$ is the time of wake (i.e. sleep offset). Whereas during sleep,

$$V_s = \nu_{sh}H - \nu_{sc}C(T) - A_s, \tag{S11}$$

$$V_w = -\nu_{ws}\tilde{Q}_{\max} + D_w, \tag{S12}$$

$$H = H(t_{\text{on}})\mathrm{e}^{(T_{\text{on}} - T)/\chi}, \tag{S13}$$

where $T_{\text{on}}$ is the time of sleep onset. Transitions between wake and sleep when $V_s = \hat{\theta}_S$, so the switch from wake to sleep occurs when

$$H \equiv H^+ = \frac{\hat{\theta}_S + A_s + \nu_{sw}\tilde{Q}_{\max} + \nu_{sc}C(T)}{\nu_{sh}}, \tag{S14}$$

and from sleep to wake when

$$H \equiv H^- = \frac{\hat{\theta}_S + A_s + \nu_{sc}C(T)}{\nu_{sh}}. \tag{S15}$$

Comparing equations (S10), (S13), (S14) and (S15) with their counterparts in the HCL model, namely

$$H(t) = \begin{cases} \mu + (H(t_{\text{off}}) - \mu)\exp\left(-\frac{t - t_{\text{off}}}{\chi}\right), & \text{during wake,} \\ H(t_{\text{on}})\exp\left(-\frac{t - t_{\text{on}}}{\chi}\right), & \text{during sleep,} \end{cases} \tag{S16}$$

and

$$\begin{aligned} H^+(t) &= \overline{H}_0 + \frac{1}{2}\Delta + c_a C(t), \\ H^-(t) &= \overline{H}_0 - \frac{1}{2}\Delta + c_a C(t), \end{aligned}$$

we see that the PR hard switch model on the slow manifold and the HCL model are equivalent if

$$\begin{aligned}\mu &= \bar{\mu} Q_m, \\ c_a &= \frac{\nu_{sc}}{\nu_{sh}}, \\ \overline{H}_0 &= \frac{\hat{\theta}_S + A_s}{\nu_{sh}} + \frac{1}{2}\frac{\nu_{sw}\tilde{Q}_{\max}}{\nu_{sh}}, \\ \Delta &= \frac{\nu_{sw} Q_m}{\nu_{sh}}.\end{aligned}$$

Hence, the HCL model is a representation of the PR model on the slow manifold for a firing function which is a hard switch.

##### A.2 Soft switch slow manifold approximation and equivalence to the HCL model

Using a similar idea, one can take the original PR model with the soft switch and on the slow manifold find expressions which relate the original PR model to the HCL model, see [1], for further details.

#### B Derivation of threshold

In the HCL model the thresholds capture wake propensity. Rather than the profile used in the original two process model, we derive a profile from measurements of wake propensity from a forced desynchrony experiment [3]. Data are reproduced in Fig. A.

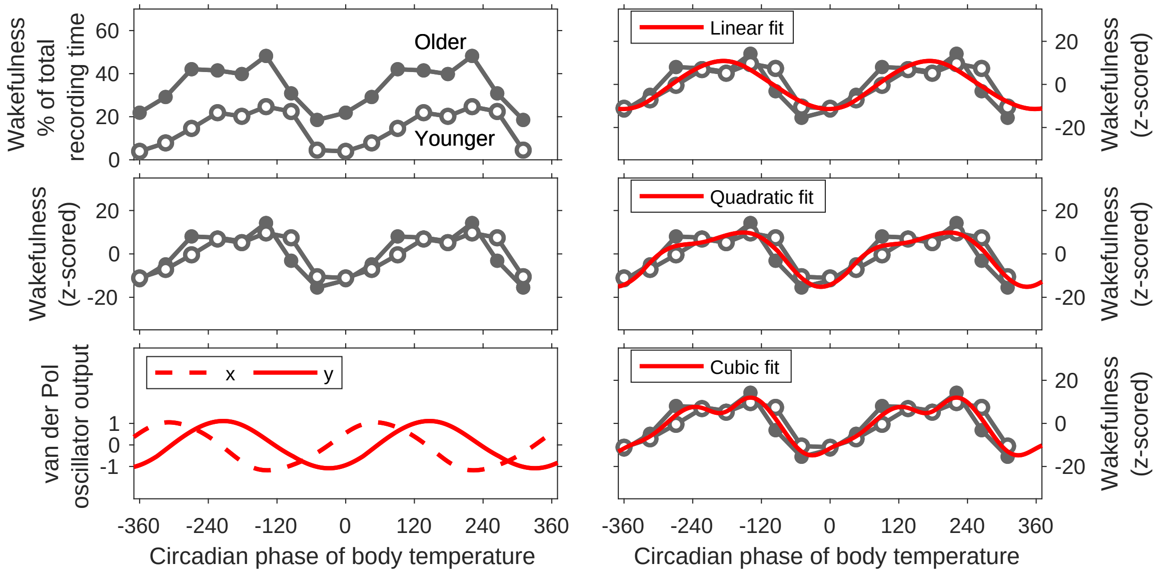


Figure A: **Derivation of the circadian drive for wakefulness by fitting to data.** In the top left panel data for the circadian rhythm of wakefulness are reproduced from [3]. Data for each group of participants were z-scored to remove the difference in mean level (central left panel). The bottom left panel shows the output variables $x$ and $y$ from the van der Pol oscillator. The right hand panels show the results of fitting to linear, quadratic and cubic functions of $x$ and $y$.

It is notable that the mean threshold levels between younger and older participants differ substantially, but that the peak-to-peak amplitude in the circadian oscillation of wakefulness is similar. In [4], changes in sleep duration across the lifespan were modelled as a change in the drive to be awake, equivalent to changing the upper asymptote, $\mu$, in the two process model. Similar results could have been obtained by changing the drive to be awake through changing $A_v$ in the neuronal models, equivalent to changing the mean threshold level $\overline{H}_0$ in the HCL model.

In order to fit the wakefulness profile, the following process was carried out. Using a default light profile, the van der Pol oscillator equations (equations (5) to (10) in the main manuscript) were integrated yielding values for $x$ and $y$. The variable $y$ was aligned with the core body temperature rhythm minus 2 hours. Then three possible $C(t)$ were constructed, namely

$$\begin{aligned} C_1(t) &= c_{10} + \alpha_{11}x + \alpha_{12}y \\ C_2(t) &= c_{20} + \alpha_{21}x + \alpha_{22}y + \beta_{21}x^2 + \beta_{22}xy + \beta_{23}y^2 \\ C_3(t) &= c_{30} + \alpha_{31}x + \alpha_{32}y + \beta_{31}x^2 + \beta_{32}xy + \beta_{33}y^2 + \gamma_{31}x^3 + \gamma_{32}x^2y + \gamma_{33}xy^2 + \gamma_{34}y^3. \end{aligned}$$

Each function was fitted to the data using a least squares minimization routine (fminunc in MATLAB). The amplitude of the resulting rhythm was then scaled to have a peak-to-peak amplitude of two i.e. oscillating from approximately -1 to 1.

| Circadian modulation of wakefulness parameters | | |
|---|---|---|
| $c_{20}$ = 0.7896 | $\alpha_{21}$ = -0.3912 | $\alpha_{22}$ = 0.7583 |
| $\beta_{21}$ = -0.4442 | $\beta_{22}$ = 0.0250 | $\beta_{23}$ = -0.9647 |

Table A: **Threshold parameter values**

We discounted the cubic function because it did not give a significantly better fit to the data ($p = 0.34$ when compared with the linear function, $p = 0.54$ when compared with the quadratic function). Although the quadratic function was not a significantly better fit to the data than the linear function ($p = 0.14$), there is considerable evidence from forced desynchrony protocols that the wakefulness profile is skewed and we therefore selected the quadratic function.

### C Comparison of the parameter regime of the HCL model with that of the original two process model

An important additional feature of the HCL model as compared with the original two process model is that it includes the process whereby light entrains the circadian rhythm. Including a process for light allows one to model the timing of sleep relative to clock time and not only relative to the timing of the circadian rhythm.

As discussed in Section A, the HCL model may be considered as a slow manifold version of a neuronal model. For concordance with previous work using a neuronal model in [5], which is an adapted version of [6], we have selected values for the homeostatic sleep constants, upper asymptote and thresholds which are consistent with neuronal models. Consequently, for the homeostatic time constant, we have elected to use the same time constant during wake and sleep of 45 hours, whereas in the two process model the homeostatic time constant is derived from slow wave activity and during wake is normally taken to be approximately 18 hours while during sleep is approximately 4 hours [7]. However, a direct comparison of the time constants and the consequent meaning for sleep homeostasis is challenging because it is not only the time constants but also the upper (and lower) asymptotes that dictate the time course. We note that the ratio of the average rate of change of sleep homeostasis during wake compared with the average rate of change of homeostasis during sleep is given by the length of wake versus the length of sleep, as can be seen as follows.

Suppose the length of wake is $\tau_w$ and sleep homeostasis takes the value $H_{\text{off}}$ at the start of the wake period and $H_{\text{on}}$ at the end of the wake period. Suppose the length of sleep is $\tau_s$. Then the average rate of change of sleep homeostasis during wake, $R_w$ is

$$R_w = \frac{H_{\text{on}} - H_{\text{off}}}{\tau_w}.$$

Similarly, the average rate of change of sleep homeostatis during sleep, $R_s$ is

$$R_s = \frac{H_{\text{off}} - H_{\text{on}}}{\tau_s}.$$

Hence,

$$\frac{R_w}{R_s} = \frac{\tau_s}{\tau_w},$$

and provided a model can simulate a fixed sleep to wake ratio, the ratio of average rise rate to decay rate will be the same, regardless of the values of the time constants. Differences between the time constant regimes will, however be more apparent if comparing, for example, the first half of the sleep episode to the second half of the sleep episode (for a decay rate of 45 hours, there is little difference between the first half and the second half of the night).

In order to further highlight the differences between these two parameter regimes, for the moment we ignore the role of light and fix the circadian input as $C(t) = \cos(\omega t)$.

In the absence of a circadian rhythm, the original two process parameters lead to a sleep-wake cycle which oscillates with a natural period $T_{\text{nat}}$ of approximately 22.5 hours (see Fig. B). Introducing a small amplitude 24-hour circadian oscillation of the thresholds then results in entrained sleep-wake cycles of 24 hours. In contrast, using time constants of 45 hours for homeostatic sleep pressure during both sleep and wake yields sleep-wake oscillations with a natural period of only approximately 10 hours in the absence of a circadian rhythm. In order to produce entrained 24-hour sleep-wake cycles a large amplitude circadian rhythm is then required. The large amplitude circadian rhythm then leads to further differences between the two-process model parameter regime and the HCL parameter regime for modelling the impact of social constraints, see below.

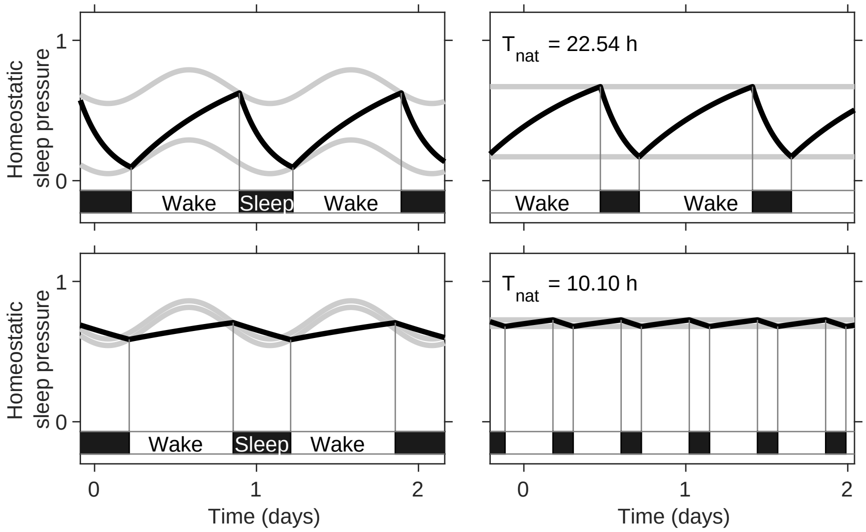


Figure B: **Comparison of the parameters for the circadian amplitude and homeostatic time constants for the original two-process model with the Homeostatic-Circadian-Light (HCL) model.** Left hand top panel shows a simulation using typical values for the original two-process model as compared with typical values for the HCL model. For ease of comparison, the HCL model has been scaled in the same way as the two-process model so that the upper asymptote is one. In addition, since the two-parameter model does not include a circadian-light interaction, the thresholds are here represented by sinusoids. In order to further highlight the differences in the homeostatic parameters, simulations for a zero circadian amplitude are shown in the right hand panels for the two different parameter regimes. The natural period of the sleep-wake cycle $T_{\mathrm{nat}}$ is much shorter for the HCL model.

### D Light metrics

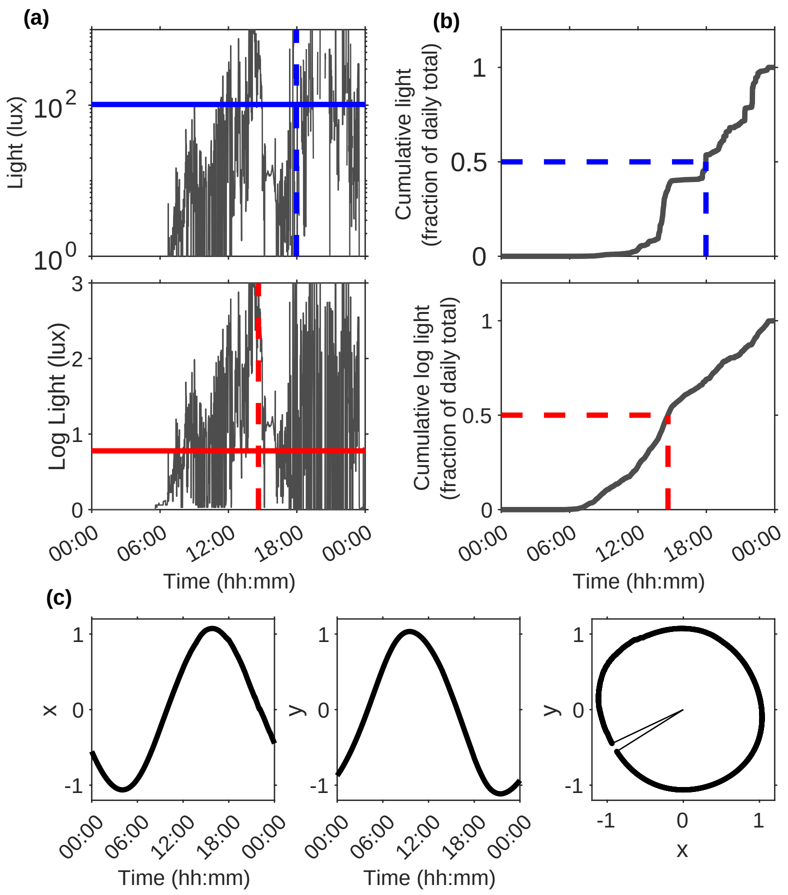


Figure C: **Light metrics.** (a) Lux levels for one day for one participant plotted on a log scale in the top panel and log (lux+1) in the bottom panel. The horizontal blue line marks the average lux value. The horizontal red line marks the average log (lux+1) value. If the average log (lux+1) is $m$, then the geometric mean is given by $10^m$. The vertical dashed lines mark the time of half light and time of half log light respectively. As shown in panels (b), these are calculated as the time at which half the daily cumulative value is reached for each day. Here, we have taken the day to run from midnight to midnight, but for populations who typically go to bed substantially after midnight calculating the cumulative time from, for example 04:00, may be more appropriate. (c) When light is fed into the forced van der Pol oscillator, it affects the time course of the van der Pol oscillator variables $x$ and $y$. After integrating for 24 hours, the circadian oscillator will have either completed exactly one cycle, less than one cycle (as shown in the far right panel of (c)) or more than one cycle. In this example, in 24 hours, the circadian clock has only completed 98.13% of a cycle, so is 27 minutes slow. In our new light metric this is recorded as -27.

### E Parameter sensitivity for light parameters

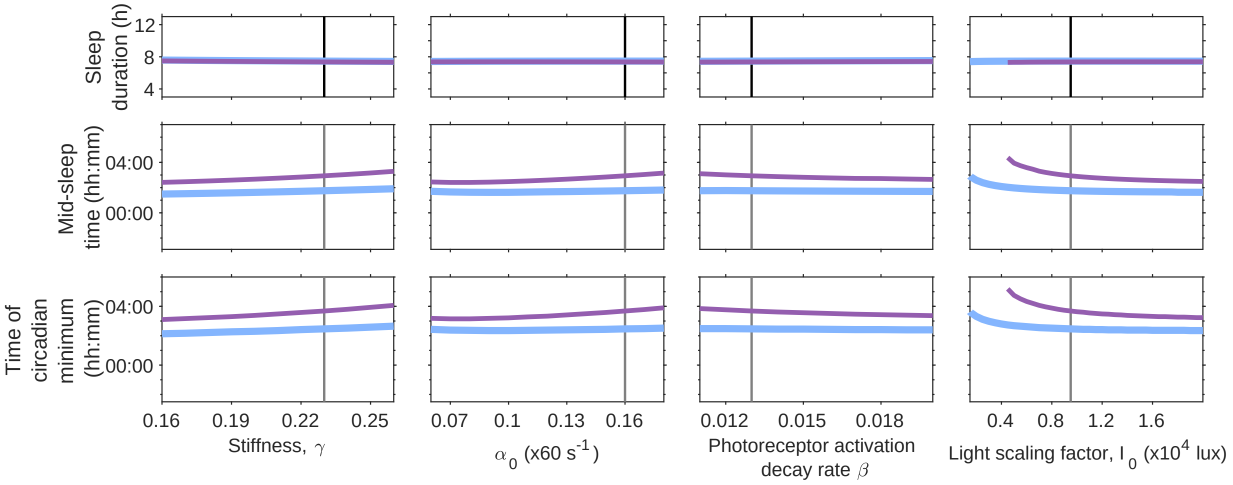


Figure D: **Dependence of model predicted sleep duration and timing on model parameters and light exposure.** In each group of panels, the dependence of sleep duration (top panels), mid-sleep time (middle panels) and time of the circadian minimum (bottom panels) for the two light availability profiles given in Fig. 2 are shown. The black vertical line indicates the default parameter values. The equivalent graphs for the five sleep-wake parameters and the remaining five circadian-light parameters are shown in Fig. 2.

### F Phase angle between the time of mid-sleep and the circadian minimum

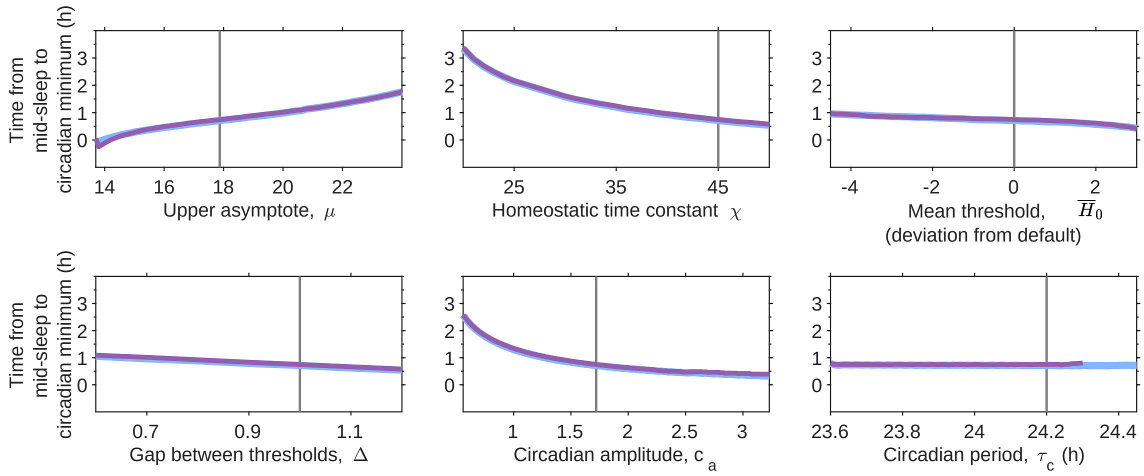


Figure E: **Phase angle between mid-sleep and the circadian minimum.** Each panel shows the number of hours from mid-sleep to the circadian minimum for the five sleep-wake parameters and the circadian period $\tau_c$ for the two light availability profiles given in Fig. 2. The black vertical line indicates the default parameter values. Like $\tau_c$, none of the other circadian-light parameters alter the phase relationship between mid-sleep and the circadian minimum so they are not shown.

#### G Sleep onset and offset

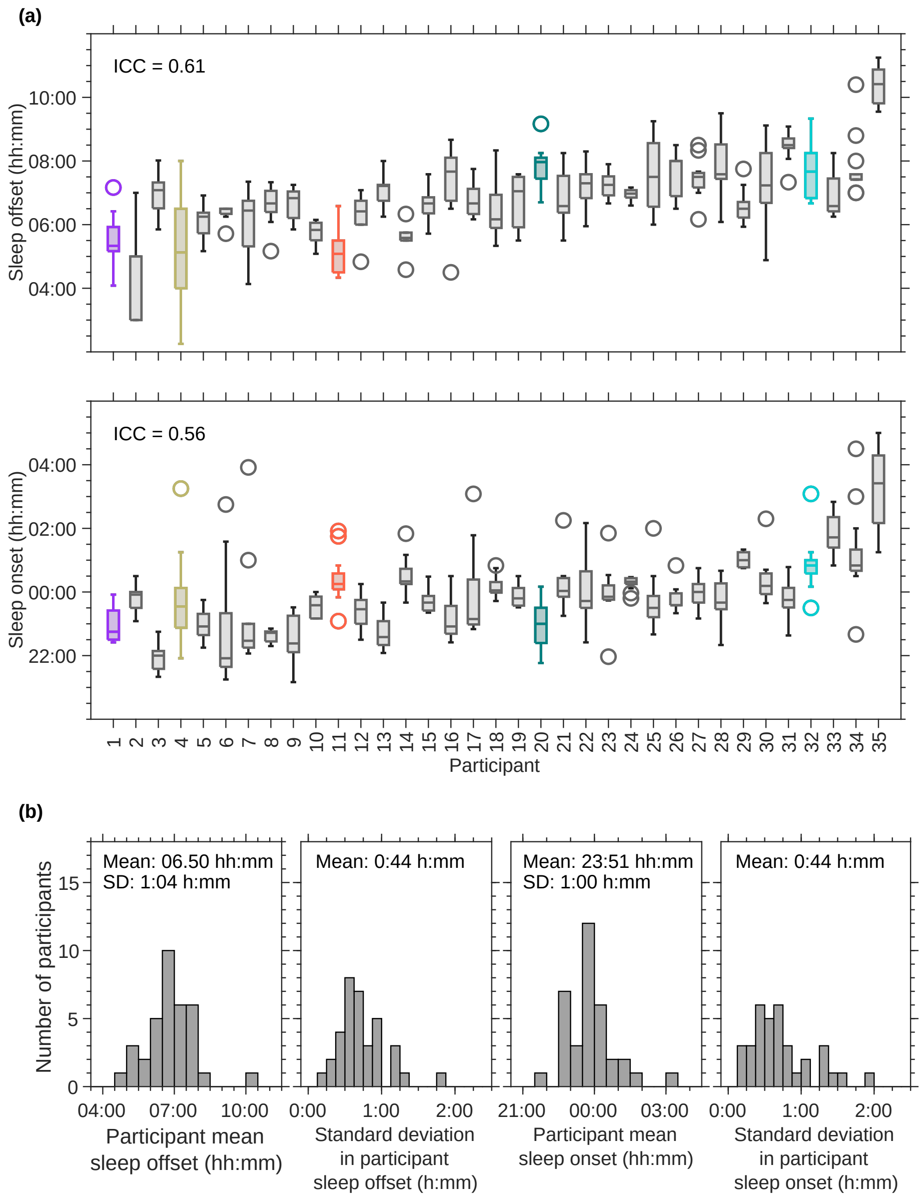


Figure F: **Summary information for sleep onset and offset for each participant and the cohort.** Top and centre panels show box plots for sleep offset and onset respectively for each participant. Bottom panels show the distributions of mean participant sleep offset, standard deviation of participant sleep offset, mean participant sleep onset and standard deviation of participant sleep onset.

### H Accuracy and correlation of fitted parameters

Fitted parameters replicate sleep duration and timing to a high accuracy, where the accuracy is essentially set by the tolerances placed on the optimization algorithm. Here, the residual $E$ for each participant is calculated as

$$E = \sqrt{\left(\overline{SD}_{\mathrm{obs}} - \overline{SD}_{\mathrm{fit}}\right)^2 + \left(\overline{MS}_{\mathrm{obs}} - \overline{MS}_{\mathrm{fit}}\right)^2}. \tag{S17}$$

where $\overline{SD}_{\mathrm{obs}}$ is the participant mean observed sleep duration, $\overline{SD}_{\mathrm{fit}}$ is the participant mean fitted sleep duration, $\overline{MS}_{\mathrm{obs}}$ is the participant mean observed time of mid-sleep and $\overline{MS}_{\mathrm{fit}}$ is the participant mean fitted time of mid-sleep.

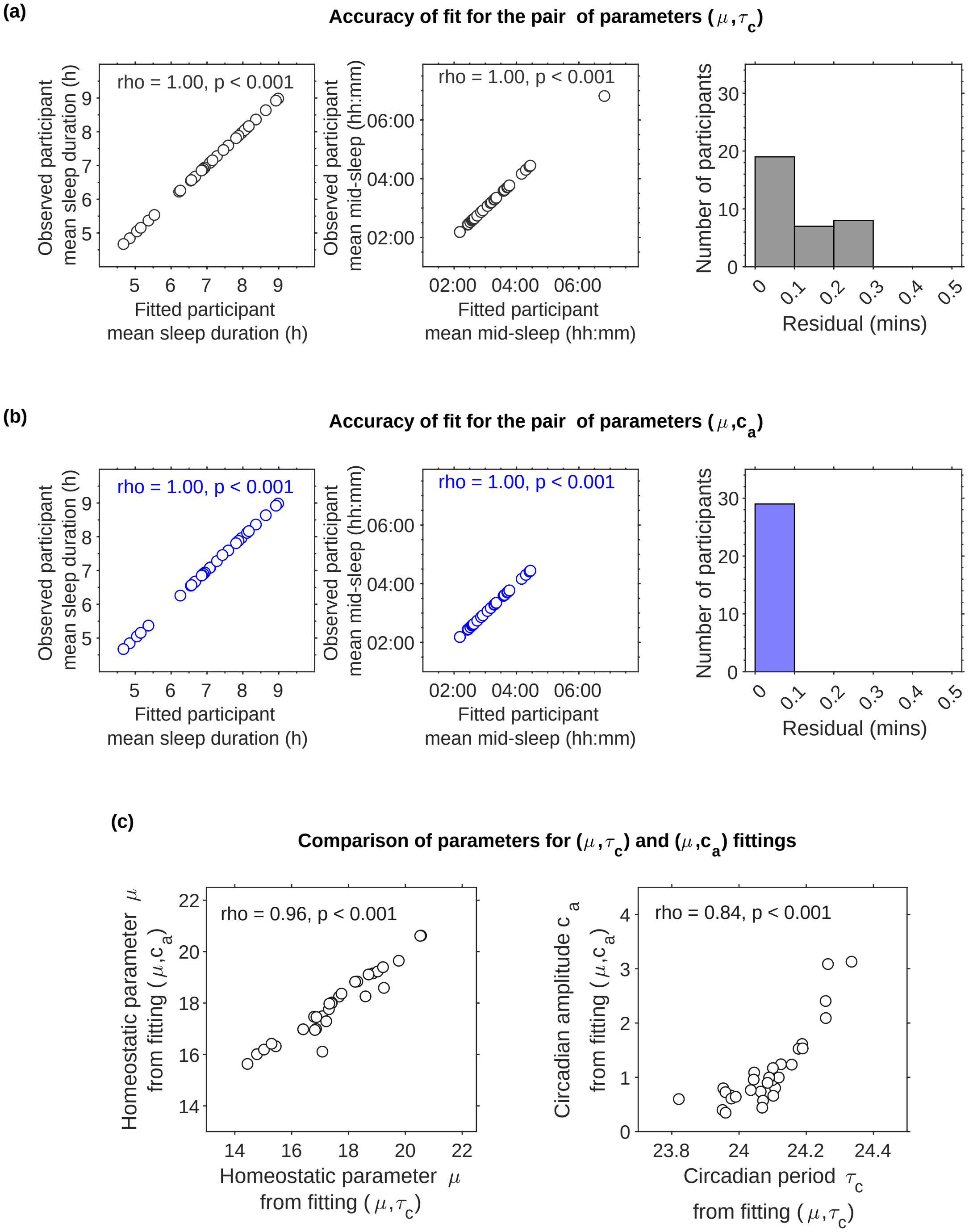


Figure G: **Accuracy and correlation of fitted parameters.** Fitted parameters accurately predict participant average sleep duration and mid-sleep timing, as shown by the very high correlation between observed and fitted values and the size of the residuals, shown in panels (a) for the pair of parameters $(\mu, \tau_c)$ are fitted and in (b) for the pair of parameters $(\mu, c_a)$. The residuals were calculated using equation (S17).

### I Sensitivity of the fitted parameters to position of the light sensor and to data imputation

Values for light exposure recorded at the wrist were typically lower than those recorded at the same time at the shoulder, see Fig. H. Prior to use, all devices were tested against a gold standard light measuring device. Nevertheless, whether attributable to differences between devices or device position, correlation between $(\mu, \tau_c)$ values calculated from light data recorded at the wrist versus light data recorded at the shoulder was extremely high, see Fig. H. The correlation for the homeostatic parameter $\mu$ of 1.00 (n=16), is primarily because the value of $\mu$ is set by sleep duration and not by the light exposure pattern. The correlation of 0.81 (n=16) between the values for $\tau_c$ is perhaps more surprising, but highlights that it is the balance of light and dark across the 24-hour day that matters, so results are relatively insensitive to factors that scale *all* light measurements up or down without change of the time and duration of the dark period.

The value of $\mu$ was insensitive to whether imputed or raw light were used. Although there was a very high correlation for $\tau_c$, differences between the first group (n=18) and second group (n=16) of the study were evident. Data were collected from the first group of participants in the late winter / early spring. Although participants were instructed to keep the wrist light monitor uncovered, on inspection of the data it was evident that there was a high percentage (41%) of zero light recordings when participants self-reported being awake, and that those zero values were distributed mainly during the evening hours. In these cases, the low levels of recorded light in the evening leads to earlier predictions of sleep timing for raw than imputed data, and consequently biasing estimates of $\tau_c$ for raw data to longer $\tau_c$, see Fig. H.

Fitting for the pair of parameters $(\mu, c_a)$ were also relatively insensitive to position of sensor and to data imputation, see Fig. I.

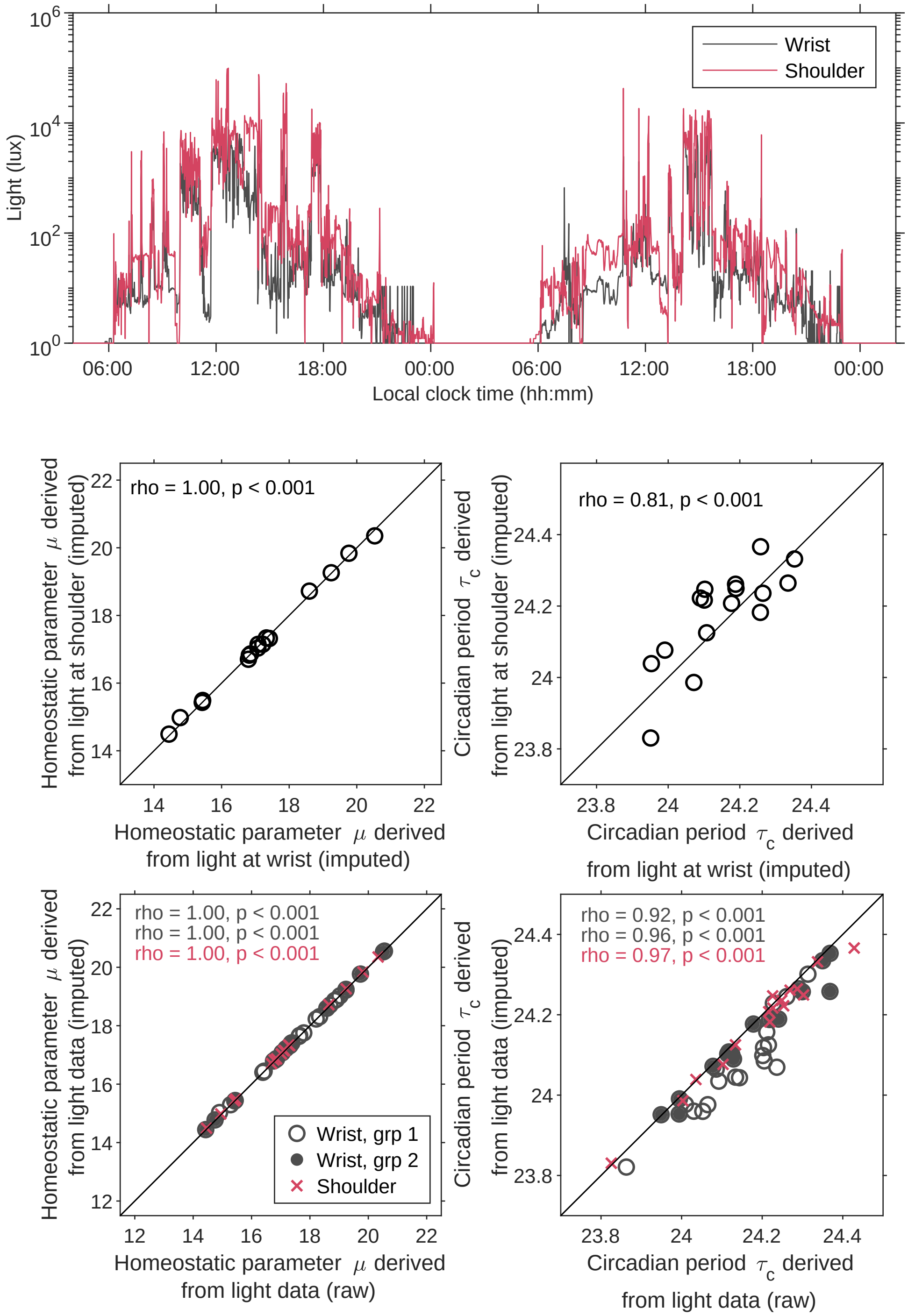


Figure H: **Sensitivity of fitted parameters $(\mu, \tau_c)$ to position of the light sensor and imputation of light data.** The top panel shows two representative days of light exposure data, collected simultaneously from the wrist and the shoulder from one participant. The central two panels show the correlation between parameters found by fitting using light collected from the wrist versus light collected from the shoulder. The lower two panels show the correlation between parameters found by fitting using raw light data versus those using imputed light data. Here, all correlations are a result of fitting for the pair of parameters $(\mu, \tau_c)$ . The equivalent correlations for the pair of parameters $(\mu, c_a)$ are shown in Fig. I.

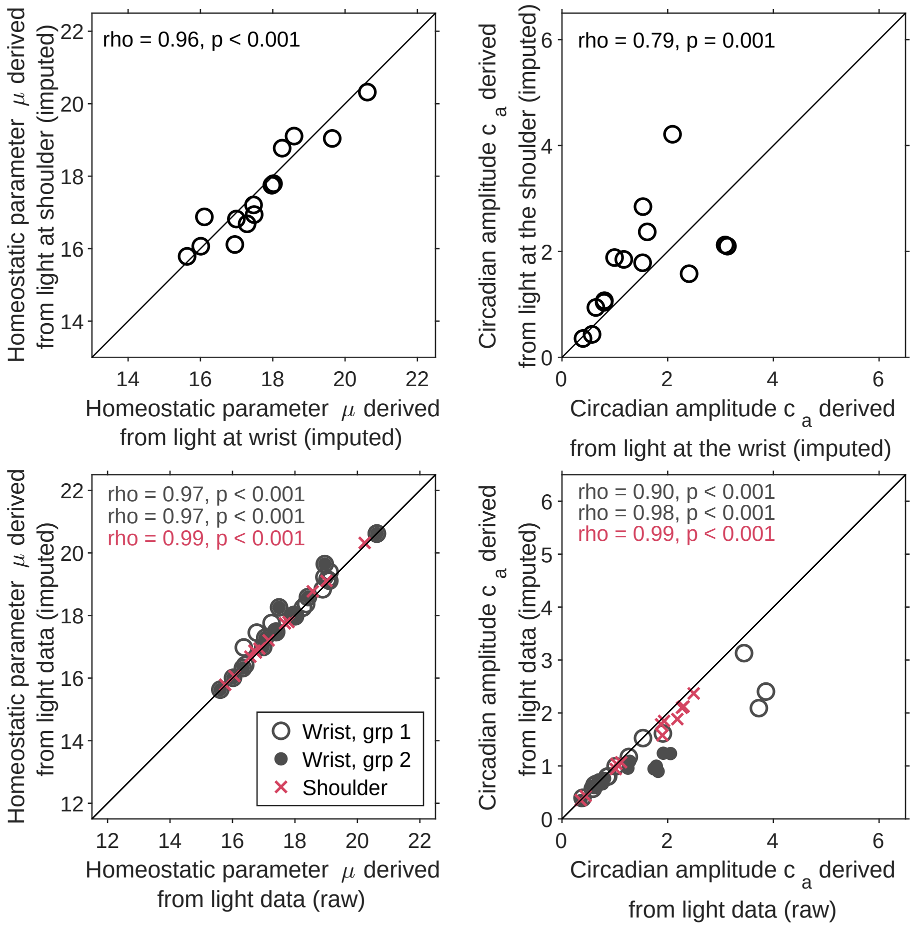


Figure I: **Sensitivity of fitted parameters $(\mu, c_a)$ to position of the light sensor and imputation of light data.** The top two panels show the correlation between parameters found by fitting using light collected from the wrist versus light collected from the shoulder. The lower two panels show the correlation between parameters found by fitting using raw light data versus those using imputed light data. Here, all correlations are a result of fitting for the pair of parameters $(\mu, c_a)$.

#### J  Amount and timing of light

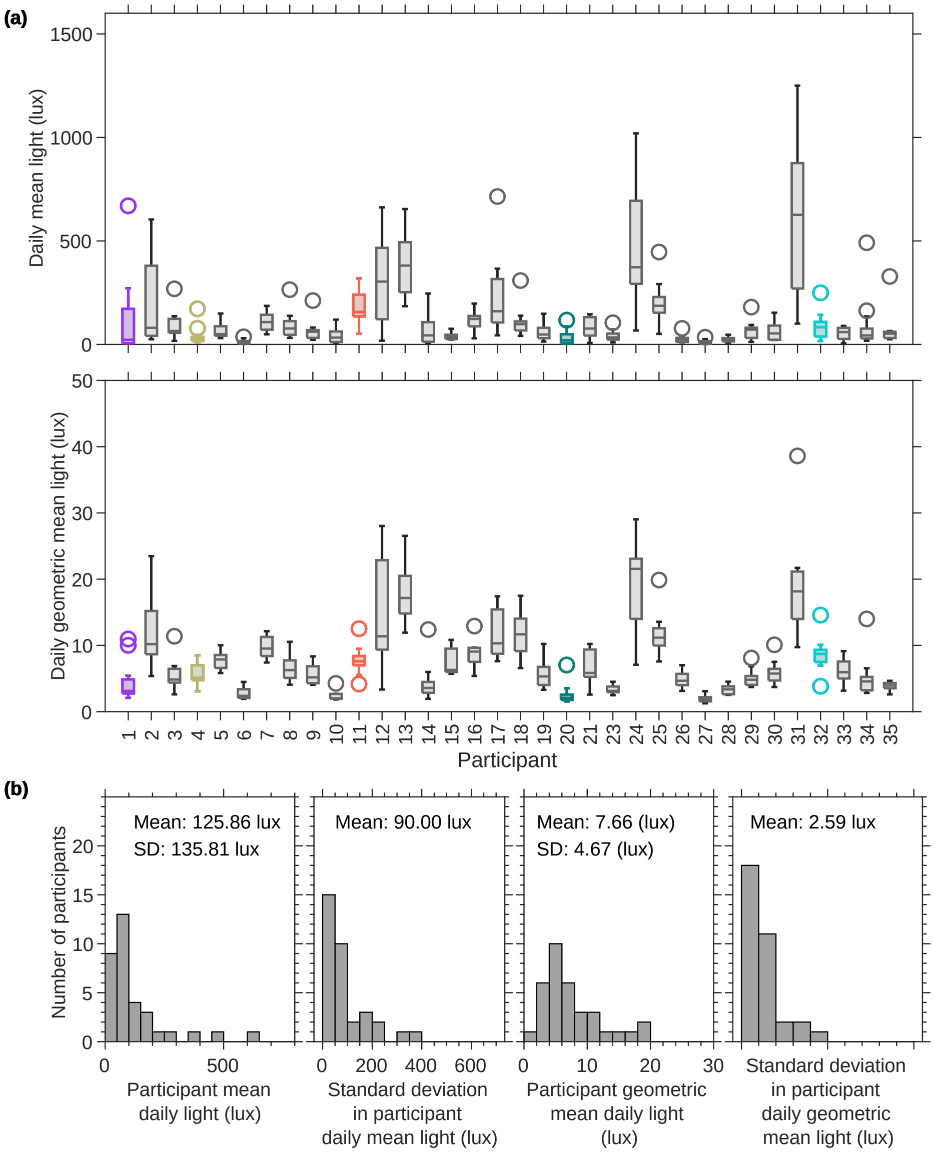


Figure J: **Summary information for measures of the amount of light exposure for each participant and the cohort.** Panels (a) show box plots for the daily mean light exposure level and the daily geometric mean of light exposure respectively for each participant. Panels (b) show the distributions of the mean and standard deviation of participant mean daily light and the mean and standard deviation of the participant geometric mean daily light.

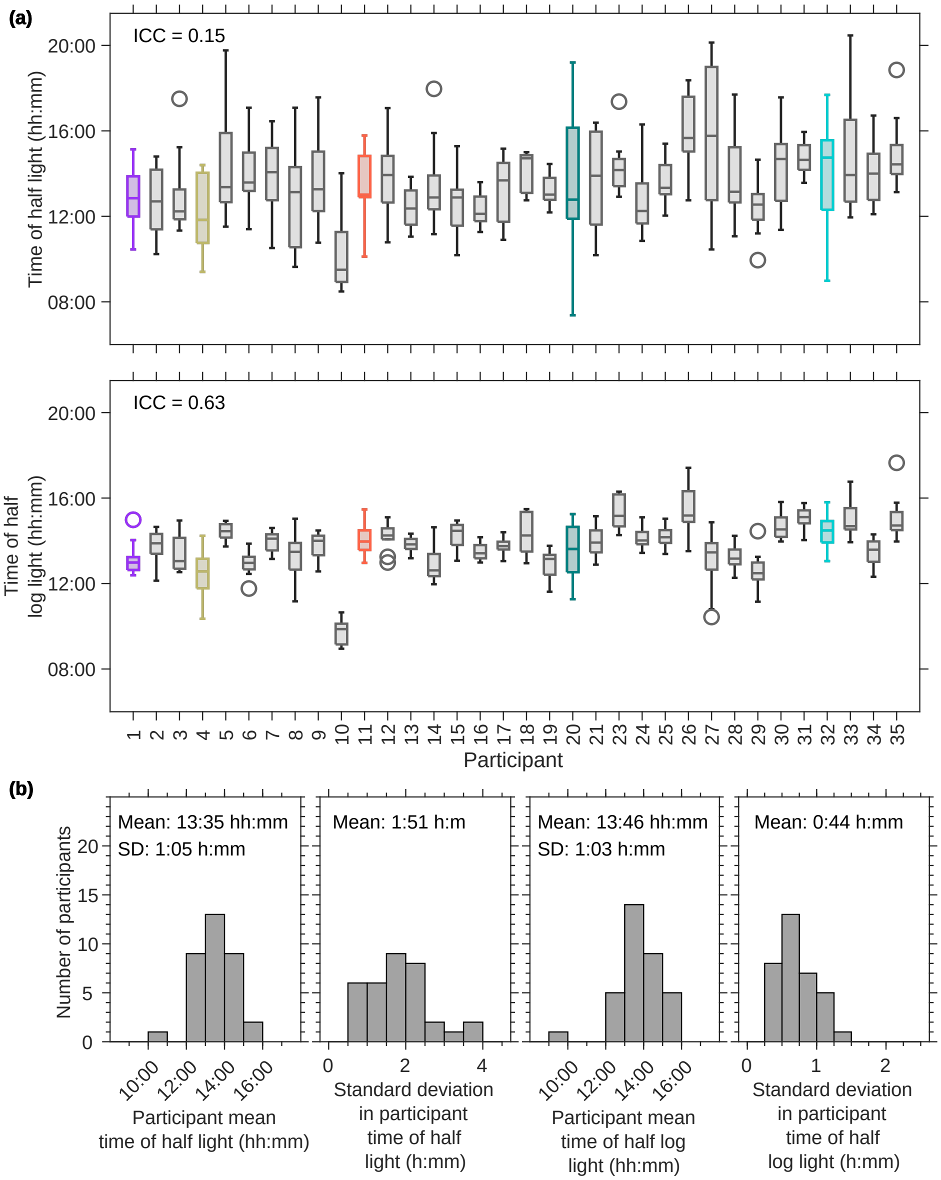


Figure K: **Summary information for measures of the timing of light exposure for each participant and the cohort.** Panels (a) show box plots for the time at which half the total daily light exposure was received in lux and log( lux +1) respectively for each participant. Panels (b) show the distributions of the mean and standard deviation of participant mean time of half light (lux) and the mean and standard deviation of the participant mean time of half light (log (lux+1)).

#### K Associations between collected data on sleep timing and duration and timing of light

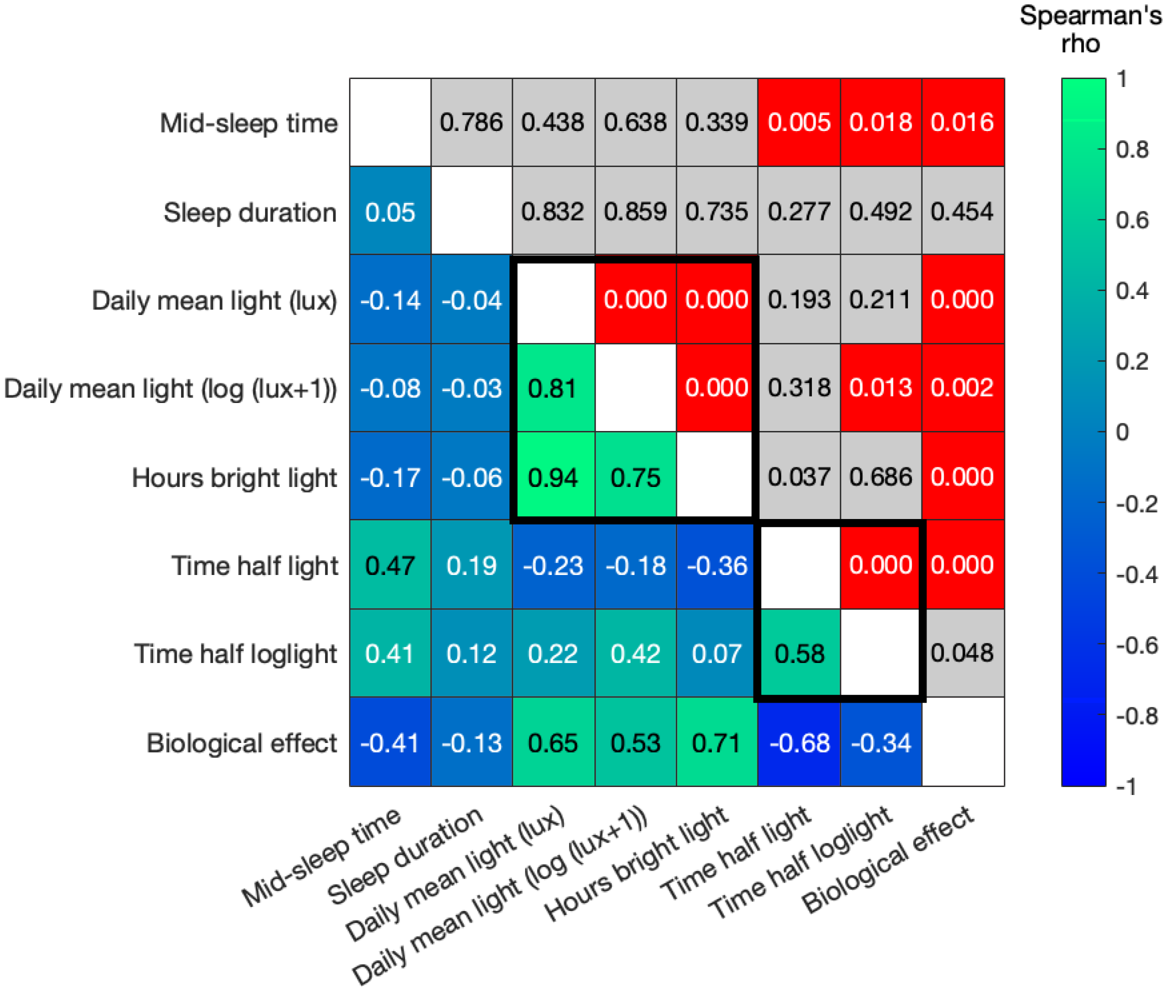


Table B: **Associations between collected data on sleep timing and duration and timing of light.** The lower triangle contains the correlation coefficients (Spearman's rho). The upper triangle indicates the p-value, shaded according to whether or not they are significant under false discovery rate correction ($p < 0.0011$). Values in grey are not significant. Associations are shown for the N=34 participants in whom light data were collected from the wrist.

#### L Entrainment tongues

Underlying the predictions of mathematical models of sleep and circadian rhythms is their formulation as forced coupled oscillator models. In these models, if the period of the forcing is too dissimilar from the intrinsic period or the 'strength' of the external forcing signal (here light) is insufficient, entrainment does not occur. Entrained regions form regions, known as Arnold tongues, on the (intrinsic period, strength of forcing) - parameter plane. For consistency with our previously published work, we constructed the HCL model so that it produces results which are similar for similar inputs. For example, in Fig. L, we show two entrainment tongues, one constructed from the simulation data used in [5] and one calculated from the HCL model. (Differences in computational speed of the HCL model compared with the neuronal model coupled with changes in the speed of computers over the last 6 years mean that we computed the HCL model with a higher accuracy, hence the sharper edges of the images).

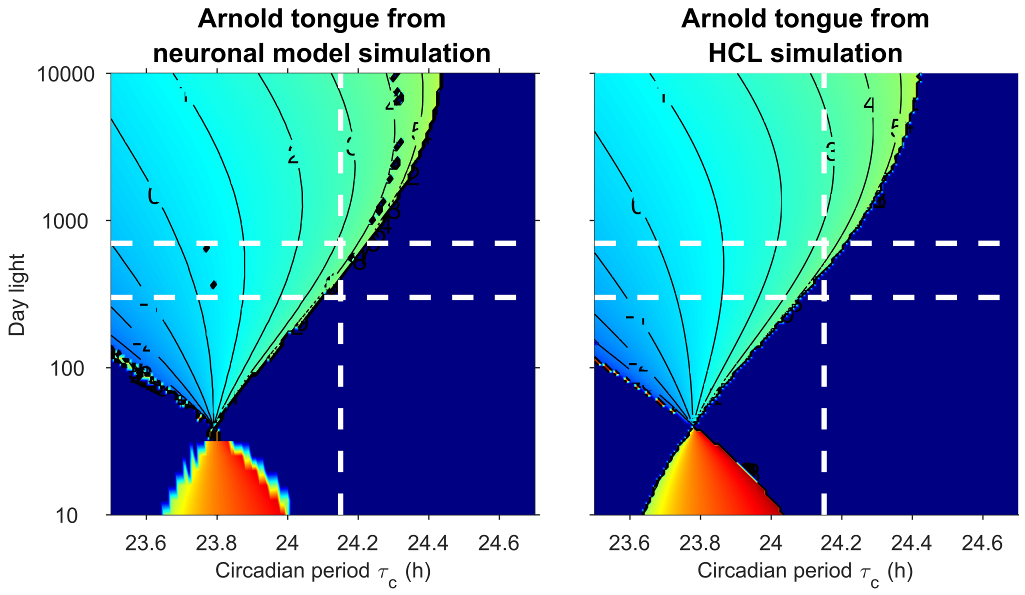


Figure L: **Entrainment tongues** for the neuronal model in [5] (left panel) and the HCL model (right panel). In both cases, the same input light profile was used – the same as the form of the light profile used for the sensitivity analysis and shown in Fig. 3 in the main text. The shading indicates mid-sleep time. The 'evening light' was set to 40. This means that when the 'day light' is also 40, the external forcing frequency takes a fixed value of 40 throughout the day and night and entrainment does not occur. For values of day light less than 40, light is brighter at night during the night than during the day and entrainment occurs to a pattern in which the sleep period is during the day (the yellow / red shaded regions).

#### M Modelling of social constraints

An attractive feature of the HCL model is the ease with which one can add additional elements in a transparent way. For example, modelling social constraints, as in [7] can be carried out by over-riding the spontaneous switching from wake to sleep and imposing wake (and / or sleep) onset times.

A similar concept for neuronal models is one of 'wake effort', i.e. that in periods when the preferred state would be sleep, that cognitive effort can enable one to override cues to sleep and remain awake. In the neuronal models, ‘wake effort’ is modelling by increasing the drive to wake promoting neurons. Within the Homeostatic-Light-Model, this concept is readily captured by defining ‘required wake’ periods that override the thresholds. The left hand panel shows an example where wake effort is used to remain awake after preferred bedtime. The right hand panel shows an example where wake effort is required to wake up before preferred wake time. We note that here too, differences between the original two-process model parameer regime and the HCL parameter regime. In the original two-process model parameter regime, during sleep, homeostasic sleep pressure remains between the two thresholds for the entire sleep period, see Fig. B. Wake effort only applies during periods when homeostatic sleep pressure is above the upper threshold, so for the original two-process parameter regime is not needed one woken early.

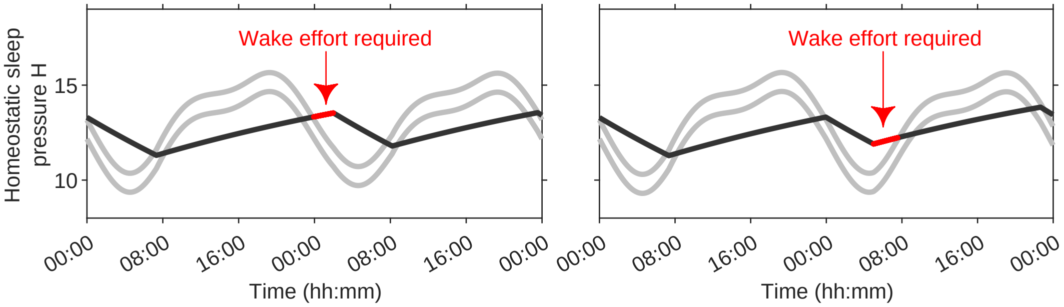


Figure M: **Modelling of social constraints and the concept of ‘wake effort’.**
